# Enterohepatic Transcription Factor CREB3L3 Protects Atherosclerosis via SREBP Competitive Inhibition

**DOI:** 10.1101/2020.04.29.069260

**Authors:** Yoshimi Nakagawa, Yunong Wang, Song-iee Han, Kanako Okuda, Asayo Oishi, Yuka Yagishita, Kae Kumagai, Hiroshi Ohno, Yoshinori Osaki, Yuhei Mizunoe, Masaya Araki, Yuki Murayama, Hitoshi Iwasaki, Morichika Konishi, Nobuyuki Itoh, Takashi Matsuzaka, Hirohito Sone, Nobuhiro Yamada, Hitoshi Shimano

## Abstract

CREB3L3 is a membrane-bound transcription factor to maintain lipid metabolism in the liver and small intestine. CREB3L3 ablation in *Ldlr*^-/-^ mice exacerbated hyperlipidemia with remnant ApoB-containing lipoprotein accumulation, developing enhanced aortic atheroma formation, whose extent was additive between liver- and intestine-specific deletion. Conversely, hepatic nuclear CREB3L3 overexpression markedly suppressed atherosclerosis with amelioration of hyperlipidemia. CREB3L3 directly upregulates anti-atherogenic FGF21 and ApoA4, whereas antagonizes hepatic SREBP-mediated lipogenic and cholesterogenic genes and regulates LXR-regulated genes involved in intestinal transport of cholesterol. CREB3L3 deficiency accumulates nuclear SREBP proteins. Because both transcriptional factors share the cleavage system for nuclear transactivation, full-length CREB3L3 and SREBPs on endoplasmic reticulum (ER) functionally inhibit each other. CREB3L3 competitively antagonizes SREBPs for ER-Golgi transport, resulting in ER retention and proteolytic activation inhibition at Golgi, and vice versa. Collectively, due to this new mechanistic interaction between CREB3L3 and SREBPs under atherogenic conditions, CREB3L3 has multi-potent protective effects against atherosclerosis.

## Introduction

cAMP responsive element-binding protein 3 like 3 (*Creb3l3*) is expressed only in liver and intestinal cells(*Lee et al., 2011*), where the CREB3L3 protein localizes in the endoplasmic reticulum (ER) and is transported to the Golgi apparatus and nucleus (*Lee et al., 2011, Omori et al., 2001, Zhang et al., 2006*). Nuclear expression of the active form of CREB3L3 in the nucleus is increased under fasting, consistent with the finding that *Creb3l3* mRNA expression is higher during fasting than refeeding (*Danno et al., 2010*). CREB3L3 reduces plasma triglyceride (TG) levels by increasing hepatic expression of apolipoprotein-encoding genes, such as apolipoprotein A4 (*Apoa4*), *Apoa5*, and *Apoc2* (*Lee et al., 2011*); these activate blood lipoprotein lipase (LPL) activity. *Creb3l3*^-/-^ mice exhibit massive hepatic lipid metabolite accumulation and significantly increased plasma TG levels, or nonalcoholic steatohepatitis when fed an atherogenic high-fat diet (*Luebke-Wheeler et al., 2008*). Apoa4 regulates HDL metabolism by activating lecithin-cholesterol acyltransferase, a key enzyme involved in cholesterol transfer to newly synthesized HDL particles (*Chen and Albers, 1985, Steinmetz and Utermann, 1985*), stimulating cholesterol efflux from macrophages (*Fournier et al., 2000*) and activating receptor-mediated uptake of HDL by hepatocytes (*Steinmetz et al., 1990*). *Apoa4*-overexpressed mice prevent atherosclerosis development (*Cohen et al., 1997, Duverger et al., 1996, Ostos et al., 2001*).

CREB3L3 and peroxisome proliferator-activated receptor alpha (PPARα) synergistically activate hepatic fibroblast growth factor 21 (FGF21) expression (*Kim et al., 2014, Nakagawa et al., 2014*). Synthesized FGF21 proteins are secreted into circulation and transported to peripheral tissues. This includes brain and skeletal muscle, as well as white adipose tissue and brown adipose tissue, in which FGF21 activates lipolysis and thermogenesis, respectively (*Fisher et al., 2012*). These effects improve diabetes and hyperlipidemia by reducing plasma glucose, insulin, TG, and cholesterol. FGF21 suppresses atherosclerotic development by reducing hypercholesterolemia, oxidative stress, and vascular smooth muscle cell proliferation via adiponectin-dependent and -independent mechanisms (*Kokkinos et al., 2017, Lin et al., 2015*). FGF21 regulates monocyte and macrophage recruitment, proliferation, and inflammatory functions in bloods and myocardial tissues, preventing macrophage accumulation, inflammation, and fibrosis (*Lin et al., 2015, Liu et al., 2013, Pan et al., 2018*).

Recently, it has been shown that CREB3L3 plays a crucial role in lipoprotein metabolism and *Ldlr*^-/-^*Creb3l3*^-/-^ mice develop significantly more atherosclerotic lesions in the aortas than *Ldlr*^-/-^ mice (*Park et al., 2016*). However, the contribution of hepatic and intestinal CREB3L3 to atherosclerosis still remains unclear. In this study, we revealed that *Ldlr*^-/-^*Creb3l3*^-/-^ mice exhibited severe atherosclerosis by upregulating sterol regulatory element-binding protein (SREBP) activation. Liver and intestine-specific CREB3L3 knockout in *Ldlr*^-/-^ mice (*Ldlr*^-/-^ LKO and *Ldlr*^-/-^ IKO) also showed accelerated atherosclerosis formation compared with *Ldlr*^-/-^ mice. Conversely, hepatic CREB3L3 overexpression (TgCREB3L3) in *Ldlr*^-/-^ (*Ldlr*^-/-^TgCREB3L3) mice suppressed atherosclerosis. Taken together, we propose that CREB3L3 in entero-hepatic circulation has a crucial role in atherosclerosis development and its mechanism is investigated.

## Results

### *Creb3l3* deletion promotes atherosclerosis with severe hyperlipidemia at an early stage of WD diet and irrespective of genders

To evaluate early stage of atherosclerosis *Ldlr*^-/-^*Creb3l3*^-/-^ mice, female and male *Ldlr*^-/-^*Creb3l3*^-/-^ mice fed a WD for 5 weeks were evaluated. Both female and male *Ldlr*^-/-^*Creb3l3*^-/-^ mice had a significant increase in atherosclerotic lesion formation in both the entire aorta and aortic root compared with control *Ldlr*^-/-^ mice, which showed barely dectable lesions at this stage (Figure 1A, 1B, Figure 1—figure supplement 1A, 1B). Plasma TG, total cholesterol (TC), and free fatty acids (FFA) levels were markedly higher in female and male *Ldlr*^-/-^*Creb3l3*^-/-^ mice than in *Ldlr*^-/-^ mice (Figure 1C, 1D, Figure 1—figure supplement 1C, 1D). High-performance liquid chromatography (HPLC) analysis revealed marked accumulation of TG and cholesterol, and significant enrichment of ApoB-containing lipoprotein fractions in female and male *Ldlr*^-/-^*Creb3l3*^-/-^ mice (Figure 1C, Figure 1—figure supplement 1C). Significant increases in very low-density lipoproteins (VLDL)-ApoB proteins (ApoB100 and ApoB40) were observed in female and male *Ldlr*^-/-^*Creb3l3*^-/-^ mice relative to *Ldlr*^-/-^ control mice (Figure 1C, Figure 1—figure supplement 1C), indicating increased numbers of both hepatic and intestinal ApoB lipoprotein particles. ApoA4, a target for CREB3L3(*Xu et al., 2014*), has been reported to be anti-atherogenic because its overexpression protects against atherosclerosis (*Cohen et al., 1997, Duverger et al., 1996, Ostos et al., 2001*). Plasma ApoA4 levels of *Ldlr*^-/-^*Creb3l3*^-/-^ mice were significantly lower than that of *Ldlr*^-/-^mice (Figure 1C, Figure 1—figure supplement 1C). Plasma levels of FGF21, anti-atherogenic hormone were significantly reduced in female and male *Ldlr*^-/-^*Creb3l3*^-/-^ mice (Figure 1D, Figure 1—figure supplement 1D). Collectively, we speculated that the absence of *Creb3l3* induced severe combined hyperlipidemia with pro-atherogenic plasma lipoprotein and hormonal profiles. It is of a note that the atherosclerosis resion was less but clearly visible in *Ldlr*^-/-^*Creb3l3*^-/-^ mice even when they consumed a normal diet, which gave rise to almost negligible atheroma in *Ldlr*^-/-^mice (Figure 1—figure supplement 2).

**Figure 1.**
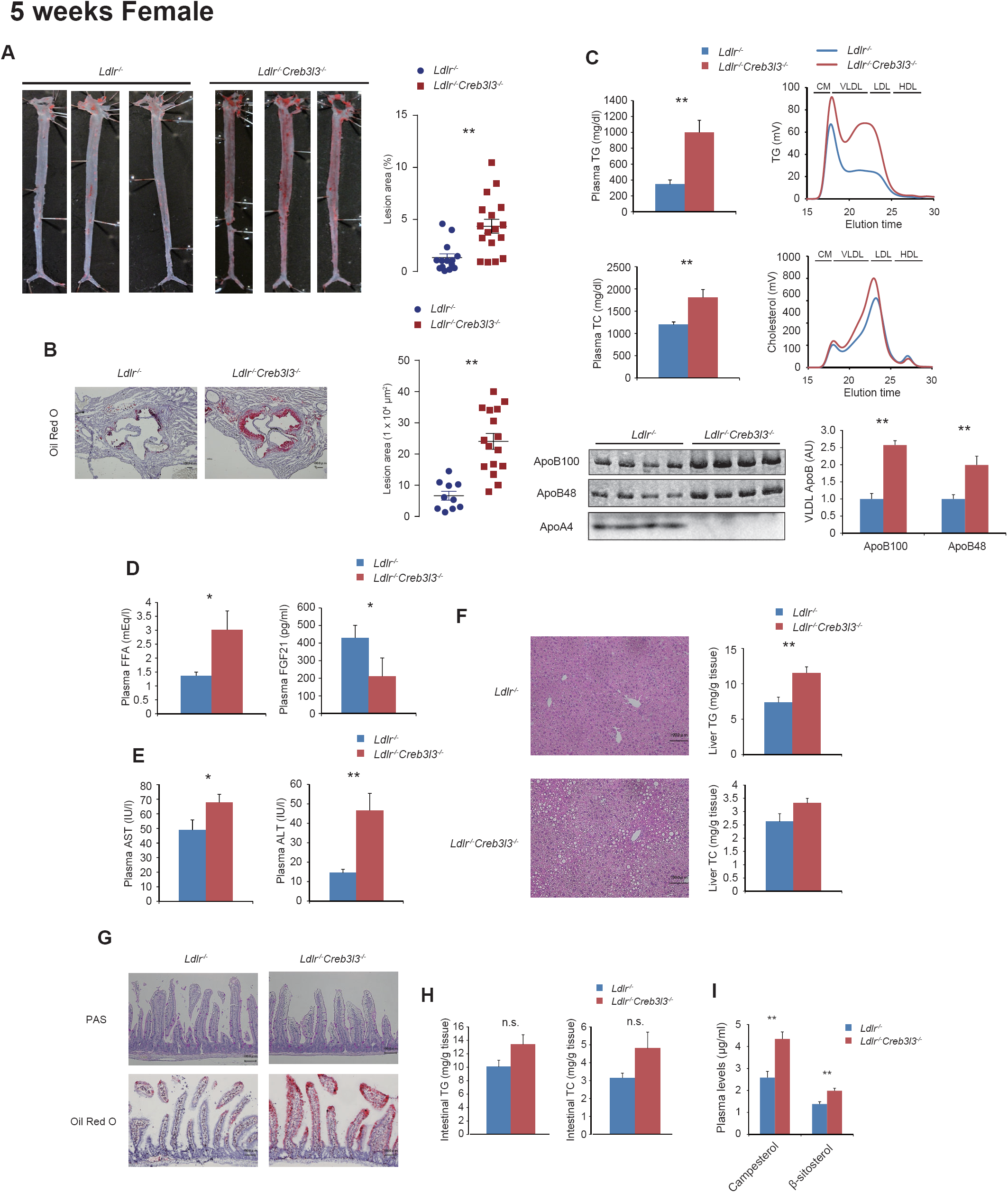
*Ldlr*^-/-^ *Creb3l3*^-/-^ mice promote atherosclerosis feeding a western diet (WD) for 5 weeks. Ten ∼ eleven-week-old female *Ldlr*^-/-^ and *Ldlr*^-/-^*Creb3l3*^-/-^ mice were fed a WD for 5 weeks; samples were collected in a fed state. (**A**) Representative images of entire Sudan IV-stained aortas from *Ldlr*^-/-^ (*n* = 14) and *Ldlr*^-/-^*Creb3l3*^-/-^ (*n* = 17) mice. The surface area occupied by lesions was quantified. \*\**p* < 0.01 vs. *Ldlr*^-/-^ mice. (**B**) Representative aortic root sections from *Ldlr*^-/-^ (*n* = 10) and *Ldlr*^-/-^*Creb3l3*^-/-^ (*n* = 16) mice. Cross-sections were stained with Oil Red O and hematoxylin. Aortic root lesion areas were quantified. \*\**p* < 0.01 vs. *Ldlr*^-/-^ mice. (**C**) Plasma TG and TC levels, *n* = 7 each; HPLC analysis of plasma lipoprotein profiles specific for plasma TG and cholesterol. ApoB100 and ApoB48 in very low-density lipoprotein (VLDL) fractions and its quantification. *n* = 7 each; \*\**p* < 0.01 vs. *Ldlr*^-/-^ mice. Plasma Apoa4 levels. **(D)** Plasma FFA (*n* = 7 each) and FGF21 levels (*n* = 5–6 per group); \**p* < 0.05 vs. *Ldlr*^-/-^ mice. **(E)** Plasma aspartate and alanine aminotransferase (AST and ALT) levels. *n* = 9–10 per group; \**p* < 0.05 and \*\**p* < 0.01 vs. *Ldlr*^-/-^ mice. (**F**) Histology of liver sections and liver TG and TC levels, *n* = 7 per group; \**p* < 0.05 and \*\**p* < 0.01 vs. *Ldlr*^-/-^ mice. (**G**) Hematoxylin and eosin staining, Oil Red O staining, and periodic acid-Schiff staining of small intestines from these mice. (**H**) Intestinal TG and TC levels of these mice. *n* = 6–8 per group (**I**) The quantification of campesterol and β-sitosterol levels of female these mice. *n* = 7–8 per group; \*\**p* < 0.01 vs. *Ldlr*^-/-^ mice.

### Deficiency of CREB3L3 dysregulates hepatic lipid metabolism and subsequently exacerbates steatohepatitis

Plasma aspartate aminotransferase (AST) and alanine aminotransferase (ALT) levels were also increased (Figure 1E), suggesting more severe liver injury in *Ldlr*^-/-^*Creb3l3*^-/-^ mice vs. *Ldlr*^-/-^ mice. Histological liver sections from female and male *Ldlr*^-/-^*Creb3l3*^-/-^ mice exhibited severe lipid accumulation (Figure 1F, Figure 1—figure supplement 1E); liver TG and TC levels in female and male *Ldlr*^-/-^*Creb3l3*^-/-^ mice were higher than those of *Ldlr*^-/-^ mice (Figure 1F, Figure 1—figure supplement 1E), supporting that *Ldlr*^-/-^*Creb3l3*^-/-^ mice increase *de novo* lipogenesis and hepatosteatosis. Taken together, we found that *Ldlr*^-/-^*Creb3l3*^-/-^ mice develop both atherosclerosis and hepatosteatosis, association of which is a recent clinical topic, regardless of gender differences. Therefore, disruption of CREB3L3 is a very strong risk factor for the onset and progression of arteriosclerosis. After this, we mainly used female mice for studying KO mice.

### Deletion of CREB3L3 in the small intestine promotes lipid absorption from diet, which contributes to hyperlipidemia

*Creb3l3* is also expressed in the intestines. Histological analysis with periodic acid-Schiff staining revealed no differences in the small intestinal mucin-producing goblet cells (Figure 1G). However, enhanced lipid accumulation in the villi of female *Ldlr*^-/-^*Creb3l3*^-/-^ mice fed a WD for 5 weeks were detected and quantitatively confirmed (Figure 1G, H), suggesting that there is a dysregulation of lipid metabolism in the small intestines of *Ldlr*^-/-^*Creb3l3*^-/-^ mice. Cholesterol absorption markers, such as campesterol and β-sitosterol levels(*Matthan et al., 2003*), in the plasma of *Ldlr*^-/-^*Creb3l3*^-/-^ mice, were significantly higher than those in *Ldlr*^-/-^ mice (Figure 1I). Taken together, the *Creb3l3* deletion in the small intestine might cause an increase in intestinal cholesterol absorption, thus exacerbating hyperlipidemia. *Ldlr*^-/-^*Creb3l3*^-/-^ mice showed an apparent increase of chylomicron production (Figure 1—figure supplement 3)., supporting that deficiency of CREB3L3 increases the activity of TG absorption in intestine and subsequent chylomicron-TG production. Taken together, CREB3L3 deletion in the small intestine contributes to hyperlipidemia. Conversely, as we previously reported, intestinal CREB3L3-overexpressed mice exhibited the suppression of plasma TC levels when fed the same diet via the suppression of cholesterol absorption in the intestine (*Kikuchi et al., 2016*). These findings indicate that hepatic CREB3L3 regulates TG metabolism, and that intestinal CREB3L3 regulates cholesterol and TG absorption in the small intestine, further suggesting that CREB3L3 regulates systemic lipid metabolism in enterohepatic circulation.

### *Ldlr*^-/-^*Creb3l3*^-/-^ mice exacerbate arteriosclerosis after a WD feeding for 3 months, a standard condition of the evaluation

*Ldlr*^-/-^*Creb3l3*^-/-^ mice exhibited early severe atherosclerosis and remained in these phenotypes even in a WD feeding for 3 months. Like when fed for 5 weeks, female *Ldlr*^-/-^*Creb3l3*^-/-^ mice revealed a significant increase in atherosclerotic lesion formation (Figure 2A, B). Plasma lipid levels were markedly higher in *Ldlr*^-/-^*Creb3l3*^-/-^ mice than in *Ldlr*^-/-^ mice (Figure 2C, D), accompanied with a marked accumulation of both TG in chylomicron, VLDL, IDL, and LDL fractions, and the entire ApoB-containing lipoproteins of *Ldlr*^-/-^*Creb3l3*^-/-^ mice (Figure 2C). Plasma FGF21 levels of *Ldlr*^-/-^*Creb3l3*^-/-^ mice were significantly lower than those of *Ldlr*^-/-^ mice (Figure 2D). These findings indicate that even when feeding a WD for 3 months, deficiency of *Creb3l3* exhibits severe atherosclerosis development with severe combined hyperlipidemia.

**Figure 2.**
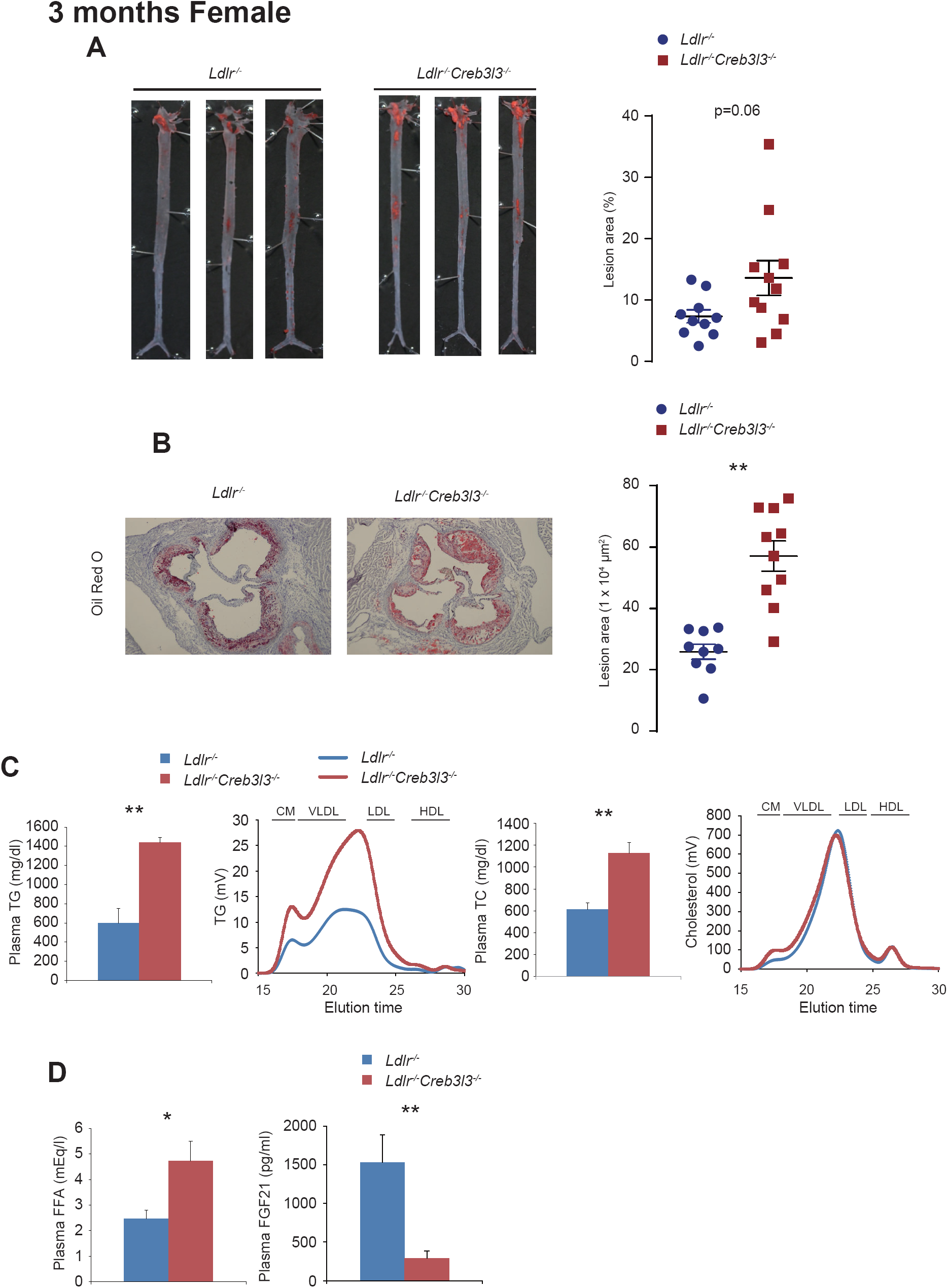
*Ldlr*^-/-^ *Creb3l3*^-/-^ mice promote atherosclerosis feeding a WD for 3 months. Ten ∼ eleven-week-old female *Ldlr*^-/-^ and *Ldlr*^-/-^*Creb3l3*^-/-^ mice were fed a WD for 3 months; samples were collected in a fed state. (**A**) Representative images of entire Sudan IV-stained aortas from *Ldlr*^-/-^ (*n* = 10) and *Ldlr*^-/-^*Creb3l3*^-/-^ (*n* = 11) mice. The surface area occupied by lesions was quantified. (**B**) Representative aortic root sections from *Ldlr*^-/-^ (*n* = 9) and *Ldlr*^-/-^*Creb3l3*^-/-^ (*n* = 10) mice. Cross-sections were stained with Oil Red O and hematoxylin. Aortic root lesion areas were quantified. \*\**p* < 0.01 vs. *Ldlr*^-/-^ mice. (**C**) Plasma TG (*n* = 6– 7))and TC (*n* = 11 each) levels. HPLC analysis of plasma lipoprotein profiles specific for plasma TG and cholesterol. \*\**p* < 0.01 vs. *Ldlr*^-/-^ mice. (**D**) Plasma FFA and FGF21 levels. *N* = 11 each; \**p* < 0.05 and \*\**p* < 0.01 vs. *Ldlr*^-/-^ mice.

### Liver and intestine CREB3L3 deficiency additively develops atherosclerosis

To define the tissue-specific contribution of CREB3L3 to suppress atherosclerosis, liver- and intestine-specific CREB3L3 knockout (LKO and IKO) mice (*Nakagawa et al., 2016a*) were crossed with *Ldlr*^-/-^ mice, generating *Ldlr*^-/-^CREB3L3 LKO and *Ldlr*^-/-^CREB3L3 IKO mice, respectively. By further crossing these mice, CREB3L3 specifically deficient in both liver and intestine of *Ldlr*^-/-^ (*Ldlr*^-/-^CREB3L3 DKO) mice were generated. The general plasma biochemical phenotypes of these mice on a normal diet were evaluated at 8 weeks; both *Ldlr*^-/-^CREB3L3 LKO and *Ldlr*^-/-^CREB3L3 IKO mice showed higher plasma TG levels than *Ldlr*^-/-^flox mice (Figure 3—figure supplement 1A). Plasma TC levels of *Ldlr*^-/-^CREB3L3 LKO were significant higher, but those of *Ldlr*^-/-^CREB3L3 IKO were not changed compared with *Ldlr*^-/-^flox mice. *Ldlr*^*-/-*^CREB3L3 DKO mice additively increased both plasma TG and TC levels additively with liver and small intestine defects (Figure S4B). HPLC analysis exhibited higher plasma TG and cholesterol levels, which were distributed over ApoB-containing lipoproteins in the order of DKO, LKO, IKO, flox mice (Figure 3—figure supplement 1A, B). DKO particularly showed a peak in the chylomicron fraction and decrease in HDL cholesterol. The pattern of plasma FFA levels was similar with that of plasma TG levels (Figure 3—figure supplement 1C). Plasma FGF21 levels of *Ldlr*^-/-^CREB3L3 LKO and *Ldlr*^-/-^CREB3L3 DKO mice were significantly lower than those of *Ldlr*^-/-^flox mice (Figure 3— figure supplement 1C), indicating that plasma FGF21 levels were dependent on hepatic CREB3L3 in contrast to both organ CREB3L3 contribution to plasma lipids. Taken together, the data indicate that both liver- and intestine-CREB3L3 additively contribute to lipid metabolism.

After feeding the mice with WD for 3 months, atherosclerotic lesion areas in either group KO mice were greater than those in control flox mice with increases in the ascending order of IKO, LKO, and DKO mice in the estimations by both the entire area and cross section at sinus (Figure 3A, B). Due to both liver and intestine absence of CREB3L3, *Ldlr*^-/-^CREB3L3 DKO mice further exacerbated atherosclerosis development, presumably by absence of both collaboratively disturbing lipid metabolism and forming atherogenic risks (Figure 3A, B).

**Figure 3.**
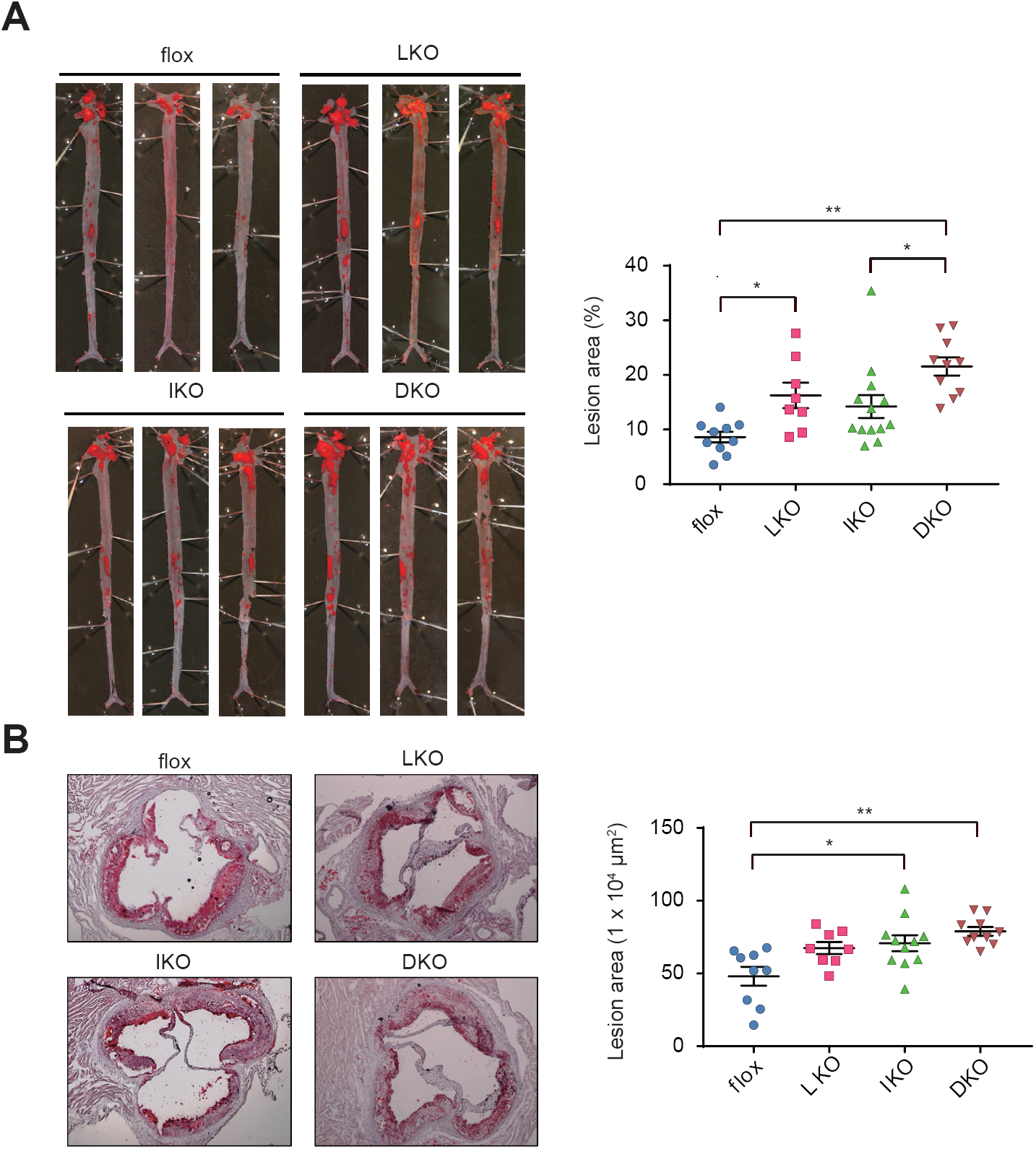
Tissue-specific *Creb3l3* deletion in *Ldlr*^-/-^ mice fed a WD for 3 months exacerbate atherosclerosis development. Ten ∼ eleven-week-old female *Ldlr*^-/-^flox (flox), *Ldlr*^-/-^liver-specific *Creb3l3* knockout (LKO), *Ldlr*^-/-^intestine-specific *Creb3l3* knockout (IKO), and *Ldlr*^-/-^liver- and intestine-specific *Creb3l3* knockout (DKO) mice were fed a WD for 3 months; samples were collected in a fed state. (**A**) Representative images of entire Sudan IV-stained aortas from flox (*n* = 10), LKO (*n* = 8), IKO, (*n* = 13) and DKO (*n* = 10) mice. The surface area occupied by the lesions was quantified. \**p* < 0.05 and \*\**p* < 0.01 among genotypes. (**B**) Representative aortic root sections from flox (*n* = 9), LKO (*n* = 8), IKO (*n* = 11), and DKO (*n* = 10) mice. Cross-sections were stained with Oil Red O and hematoxylin. Aortic root lesion areas were quantified. \**p* < 0.05 and \*\**p* < 0.01 among genotypes.

### Hepatic CREB3L3 activation suppresses formation of atherosclerotic lesions in *Ldlr*^-/-^ mice that were fed a WD

Mice with hepatic overexpression of active CREB3L3 (TgCREB3L3; Figure 4— figure supplement 1) (*Nakagawa et al., 2014*) were crossed with *Ldlr*^-/-^ mice. *Ldlr*^-/-^ and *Ldlr*^-/-^ TgCREB3L3 mice were fed a WD for 3 months and then were subjected to an atherosclerosis analysis. Lesions were markedly suppressed in both the entire aorta and aortic root of *Ldlr*^-/-^TgCREB3L3 mice (Figure 4A, B), indicating that hepatic CREB3L3 overexpression attenuates WD-induced atherosclerosis development. Additionally, because FGF21, a main CREB3L3 target, has been reported to have a protective effect against atherosclerosis, the contribution of FGF21 in these phenotypes was estimated by crossing *Ldlr*^-/-^TgCREB3L3 mice with *Fgf21*^-/-^ mice to generate *Ldlr*^-/-^TgCREB3L3*Fgf21*^-/-^ mice, followed by fed on a WD diet for 3 months. Deletion of FGF21 was confirmed by showing that plasma FGF21 levels were significantly increased in *Ldlr*^-/-^TgCREB3L3 mice and not detected in *Fgf21*^-/-^ background mice (Figure 4C). Consistent with a previous report (*Lin et al., 2015*), *Ldlr*^-/-^*Fgf21*^-/-^ mice showed a trend of more severe atherosclerosis development. However, *Ldlr*^-/-^*Fgf21*^-/-^ TgCREB3L3 mice still maintained significant suppression of atherosclerosis to similar extent in the presence of FGF21 as inhibition rates by CREB3L3 overexpression was estimated to be about 50% in *Ldlr*^-/-^ mice and about 40% in *Ldlr*^-/-^*Fgf21*^-/-^ mice (Figure 4A, B). The data suggested that anti-atherogenic effect of CREB3L3 is not mediated primarily through FGF21. CREB3L3 overexpression significantly reduced plasma TG and FFA levels of *Ldlr*^-/-^ mice and even *Ldlr*^-/-^*Fgf21*^-/-^ mice. There were no differences in plasma TC levels among all genotypes. (Figure 4C). As a causative factor for hyperlipidemia and atherosclerosis in a previous report (*Lee et al., 2011*), plasma levels of ApoA4, an LPL modulator, and a CREB3L3 target gene (*Lee et al., 2011*), were similarly increased in mice overexpressing CREB3L3 in both *Ldlr*^-/-^TgCREB3L3 and *Ldlr*^-/-^*Fgf21*^-/-^TgCREB3L3 mice (Figure 4D). In gain of function, CREB3L3 target, ApoA4, but not FGF21, could contribute to the suppressive effects of hepatic CREB3L3 on the development of atherosclerosis. Taken together with the observations from KO mice, it can be concluded that CREB3L3 prevents atherosclerosis.

**Figure 4.**
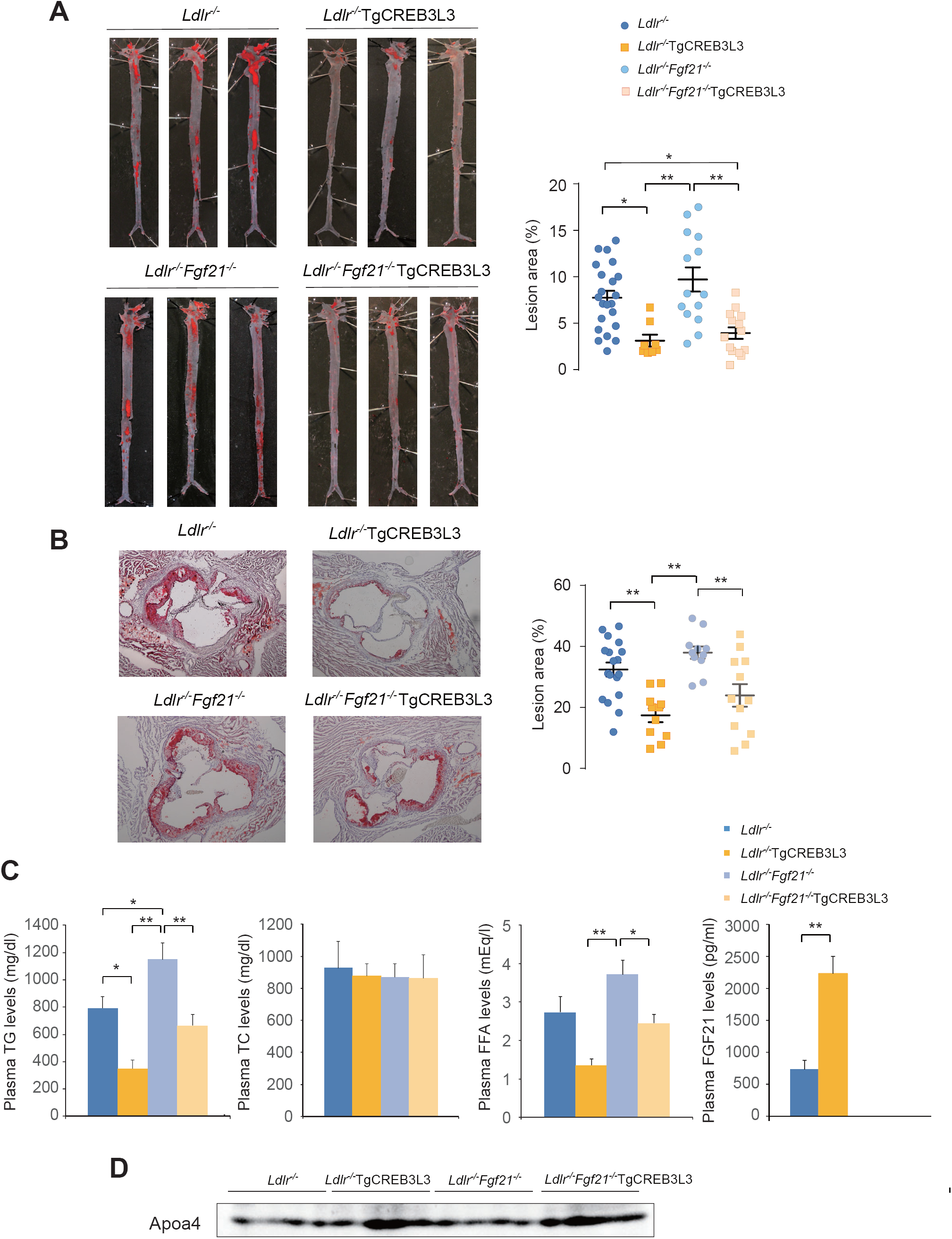
*Ldlr*^-/-^TgCREB3L3 mice suppress atherosclerosis development when fed a WD for 3 months. Ten ∼ eleven-week-old male *Ldlr*^-/-^, *Ldlr*^-/-^TgCREB3L3, *Ldlr*^-/-^*Fgf21*^-/-^, and *Ldlr*^-/-^*Fgf21*^-/-^TgCREB3L3 mice were fed a WD for 3 months. Samples were collected in a fed state. (**A**) Representative images of entire Sudan IV-stained aortas from *Ldlr*^-/-^ (*n* = 22), *Ldlr*^-/-^TgCREB3L3 (*n* = 8), *Ldlr*^-/-^*Fgf21*^-/-^ (*n* = 14), and *Ldlr*^-/-^*Fgf21*^-/-^TgCREB3L3 (*n* = 14) mice. The surface area occupied by lesions was quantified. \**p* < 0.05 and \*\**p* < 0.01 among genotypes. (**B**) Representative aortic root sections from *Ldlr*^-/-^ (*n* = 18), *Ldlr*^-/-^TgCREB3L3 (*n* = 11), *Ldlr*^-/-^*Fgf21*^-/-^ (*n* = 11), and *Ldlr*^-/-^*Fgf21*^-/-^TgCREB3L3 (*n* = 12) mice. Cross-sections were stained with Oil Red O and hematoxylin. Aortic root lesion areas were quantified. \*\**p* < 0.01 among genotypes. (**C**) Plasma TG, TC, FFA, and FGF21 levels. *n* = 7–21; \**p* < 0.05 and \*\**p* < 0.01 among genotypes. (**D**) Plasma Apoa4 levels were detected by western blotting.

### CREB3L3 regulates TG metabolism by controlling apolipoproteins in the liver of *Ldlr*^-/-^ mice

We next investigated the potential risk factors linking to atherosclerosis-prone CREB3L3 deficiency, starting with TG metabolism. VLDL secretions from the liver were significantly increased in *Ldlr*^-/-^*Creb3l3*^-/-^ mice (Figure 5—figure supplement 1A). Consistent with a previous study (*Lee et al., 2011*), the expression of LPL activators, such as *Apoc2* and *Apoa5*, was reduced significantly in *Ldlr*^-/-^*Creb3l3*^-/-^ mice, whereas the expression of the LPL inhibitor *Apoc3* was increased (Figure 5A). One of the LPL activators, *Apoa4*, tended to decrease but not significantly (Figure 5A). These changes contribute to hypertriglyceridemia by inhibiting LPL activity and impairing TG clearance. In accordance with decreased plasma LPL activity, TG clearance was decreased remarkably in *Ldlr*^-/-^*Creb3l3*^-/-^ mice (Figure 5—figure supplement 1B, C).

**Figure 5.**
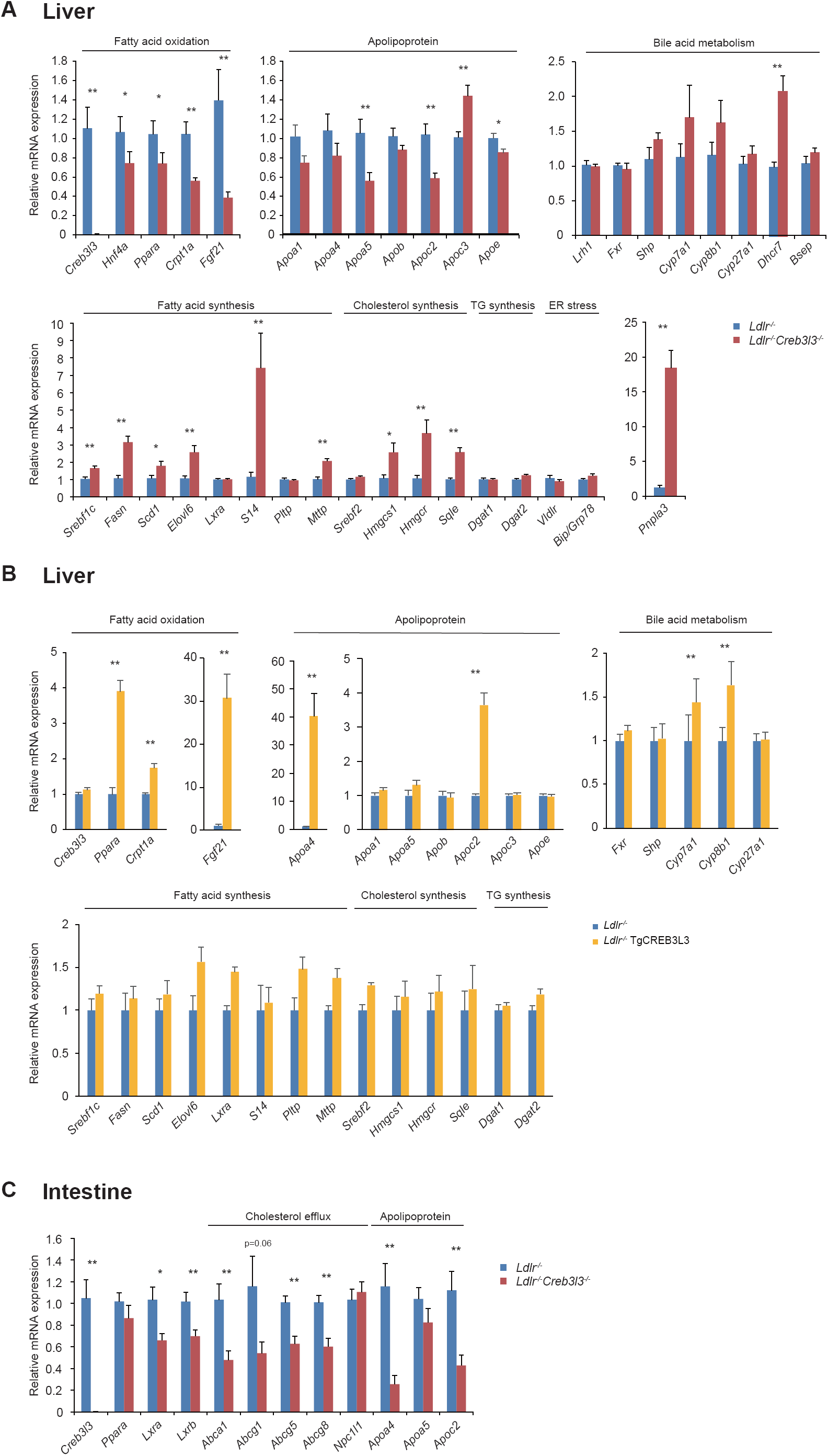
Gene expression in *Ldlr*^-/-^ mice, *Ldlr*^-/-^*Creb3l3*^-/-^ mice, and *Ldlr*^-/-^ TgCREB3L3 mice. Gene expression in the livers of 8-week-old female *Ldlr*^-/-^ and *Ldlr*^-/-^*Creb3l3*^-/-^ mice (*n* = 11 per group) (**A**) and male *Ldlr*^-/-^ and *Ldlr*^-/-^ TgCREB3L3 mice (*n* = 7 per group) (**B**) in a fed state with normal chow. \**p* < 0.05 and \*\**p* < 0.01 vs. *Ldlr*^-/-^ mice. (**C**) Gene expression in the intestines of 8-week-old male *Ldlr*^-/-^ and *Ldlr*^-/-^ TgCREB3L3 mice in a fed state with normal chow. *n* = 7 per group; \**p* < 0.05 and \*\**p* < 0.01 vs. *Ldlr*^-/-^ mice.

In contrast to *Ldlr*^-/-^*Creb3l3*^-/-^ mice, hepatic CREB3L3 overexpression significantly increased hepatic *Apoa4* and *Apoc2* expression (Figure 5B). Consistent with a previous report (*Chanda et al., 2013*), CREB3L3 overexpression increased bile acid synthesis-related gene expression, including *Cyp7a1* and *Cyp8b1* (Figure 5B). *Ldlr*^-/-^TgCREB3L3 mice exhibited an apparent increase in TG clearance (Figure 5—figure supplement 1E) but no difference in VLDL secretions compared with *Ldlr*^-/-^ mice (Figure 5—figure supplement 1D). There were no differences in plasma LPL activity between *Ldlr*^-/-^ and *Ldlr*^-/-^TgCREB3L3 mice (Figure 5—figure supplement 1F). However, changes in apolipoproteins could partially modulate plasma LPL activity and lead to consequent decrease in plasma TG-rich lipoprotein levels. FGF21 has also the ability to reduce plasma TG levels (*Schlein et al., 2016*). Our findings suggest that CREB3L3 activates *Fgf21* and LPL regulatory genes, resulting in a reduction in plasma TG levels.

### Deficiency of CREB3L3 in the small intestine of *Ldlr*^-/-^ mice dysregulates LXR signaling

Next, we focused on intestinal lipid metabolism. The expressions of *Lxra*/*b* and liver X receptor (LXR) signaling molecules, *Abca1, Abcg5*, and *Abcg8*, were significantly downregulated in the intestines of *Ldlr*^-/-^*Creb3l3*^-/-^ mice, whereas *Abcg1* tended to decrease but not significantly (Figure 5C). Based upon the previous report that intestinal overexpression of active LXRα in *Ldlr*^-/-^ (*Ldlr*^-/-^TgLXRα) mice improves atherosclerosis (*Lo Sasso et al., 2010*), we also speculated that the suppression of LXR signaling in intestines of *Ldlr*^-/-^*Creb3l3*^-/-^ mice contributed to the acceleration of atherosclerosis. Because *Ldlr*^-/-^TgLXRα mice consistently exhibited significantly reduced intestinal cholesterol absorption (*Lo Sasso et al., 2010*), the increase in intestinal cholesterol absorption in *Ldlr*^-/-^*Creb3l3*^-/-^ mice might depend on the downregulation of LXR signaling in the small intestine. Reductions in *Abcg5/8*, which increased cholesterol excretion into the intestinal lumen, also led to cholesterol accumulation in the intestines of *Ldlr*^-/-^*Creb3l3*^-/-^ mice. Similar to the liver, *Apoa4* and *Apoc2* expression were decreased in the small intestines of *Ldlr*^-/-^*Creb3l3*^-/-^ mice (Figure 5C).

### Deficiency of CREB3L3 activates hepatic expressions of SREBP-1 and −2 target genes in the liver of *Ldlr*^-/-^ mice

To further explore the integral mechanism, we determined hepatic gene expression profiles of *Creb3l3*^*-/-*^ and hepatic Tg mice (Figure 5A). Consistent with described profiles of the *Creb3l3*^-/-^ mice (*Nakagawa et al., 2014*), genes downstream of CREB3L3, including fatty acid oxidation-related genes such as *Ppara, Cpt1a* (carnitine palmitoyltransferase 1a, liver), and *Fgf21*,were decreased in *Ldlr*^-/-^*Creb3l3*^-/-^ mice. Hepatic CREB3L3 overexpression significantly increased hepatic *Ppara* as well as and its target gene expression, such as those of *Cpt1a* and *Fgf21* in *Ldlr*^-/-^ mice (Figure 5B). Lipogenic genes regulated by SREBP-1c were entirely upregulated in *Ldlr*^-/-^*Creb3l3*^-/-^ mice including *Fasn* (fatty acid synthase), *Scd1* (stearoyl-coenzyme A desaturase 1), *Elovl6* (ELOVL family member 6, elongation of long-chain fatty acids [yeast]) (Figure 5A). Another SREBP-1 target, *Pnpla3* (patatin-like phospholipase domain containing 3), which is a central regulator of hepatic TG metabolism and fat accumulation (*Huang et al., 2010*), was also remarkably increased in *Ldlr*^-/-^*Creb3l3*^-/-^ mice (Figure 5A). It was noteworthy that in contrast to marked induction of target genes, the expression of its encoding gene *Srebf1* only slightly increased. The expression of cholesterol synthesis genes governed by SREBP-2, such as *Hmgcs1* (HMGCoA synthase 1), *Hmgcr* (HMGCoA reductase), and *Sqle* (Squalene epoxidase), was increased in *Ldlr*^-/-^*Creb3l3*^-/-^ mice, although the encoding gene *Srebf2* did not exhibit apparent changes (Figure 5A). These changes in SREBP-related genes implicate the functional activation of SREBPs at the posttranslational level. In contrast, the expression of *Srebfs* and its target genes per se was not altered between *Ldlr*^-/-^ and *Ldlr*^-/-^TgCREB3L3 mice despite the hepatic overexpression of nuclear, and not full-length, CREB3L3 (Figure 5B).

### Interaction between CREB3L3 and SREBP in hepatic lipid metabolism

To explain that the hepatic expression of lipogenic and cholesterogenic genes was strongly upregulated in *Ldlr*^-/-^*Creb3l3*^-/-^ mice without appreciable induction of SREBP expression, we evaluated the proteolytic cleavage of SREBPs by the amount of precursor and nuclear SREBP proteins. Western blotting revealed that levels of both premature (membrane; pSREBP-1) and active (nuclear) forms of SREBP-1 (nSREBP-1) as well as the active form of SREBP-2 (nSREBP-2) were robustly increased in livers from *Ldlr*^-/-^*Creb3l3*^-/-^ mice (Figure 6A), which suggests activated proteocleavage of both proteins and warrants the activation of these target genes. Because SREBPs and CREB3L3 share a set of proteases (S1P and S2P) involved in the cleavage process of transcriptional activation at Golgi (*Zhang et al., 2006*), the cleavage of these proteins may be competitive to each other. Specifically, we hypothesized that the presence of premature CREB3L3 (pCREB3L3) inhibits pSREBPs cleavage in a competitive manner. Accordingly, pCREB3L3 expression was restored in *Ldlr*^-/-^*Creb3l3*^-/-^ mice via infection with adenovirus encoding pCREB3L3 (Ad-pCREB3L3) to determine whether pCREB3L3 would suppress pSREBP cleavage, thus reducing nSREBP accumulation. As expected, pCREB3L3 overexpression reduced nSREBP-1 and nSREBP-2 accumulation in *Ldlr*^-/-^*Creb3l3*^-/-^ mice with only small changes in *Srebfs* expression (Figure 6B). pCREB3L3 overexpression decreased plasma TG and TC levels compared with control GFP infected mice (Figure 6B). Genes related to SREBP cleavage were also investigated. An SREBP cleavage activator, *Scap* was increased in *Ldlr*^-/-^*Creb3l3*^-/-^ mice (Figure 6C), partly explaining that SREBPs cleavage was activated in *Ldlr*^-/-^*Creb3l3*^-/-^ mice. Consistent with the previous report, *Insig2a*, a retention factor of SREBP-SCAP complex in ER, is a target gene for CREB3L3 (*Wang et al., 2016*), *Insig2a* was decreased in *Ldlr*^-/-^*Creb3l3*^-/-^ mice (Figure 6C). Other retention factors such as *Insig1* and *Insig2b* were not changed in both *Ldlr*^-/-^*Creb3l3*^-/-^ mice (Figure 6C), meanwhile overexpression of nuclear CREB3L3 in *Ldlr*^-/-^ mice failed to affect the expression of *Insig* genes (Figure 6C). This discrepancy between loss and gain of CREB3L3 in *Ldlr*^-/-^ mice indicates that CREB3L3-Insig2a pathway is not strong enough to provide an explanation for the strong SREBP activation by CREB3L3 deficiency, supporting our new hypothesis above. In a cell-based reporter assay using an SREBP response element (SRE) containing luciferase (SRE-Luc), endogenous SREBP cleavage activity was detected by transfection with pSREBP-1a, as evidenced by SRE-Luc activity. Further co-transfection with pCREB3L3 significantly suppressed this pSREBP-1a cleavage, but the active form of CREB3L3 (nCREB3L3) did not (Figure 6D) as another supportive data for competition in the cleavage between pCREB3L3 and pSREBP.

**Figure 6.**
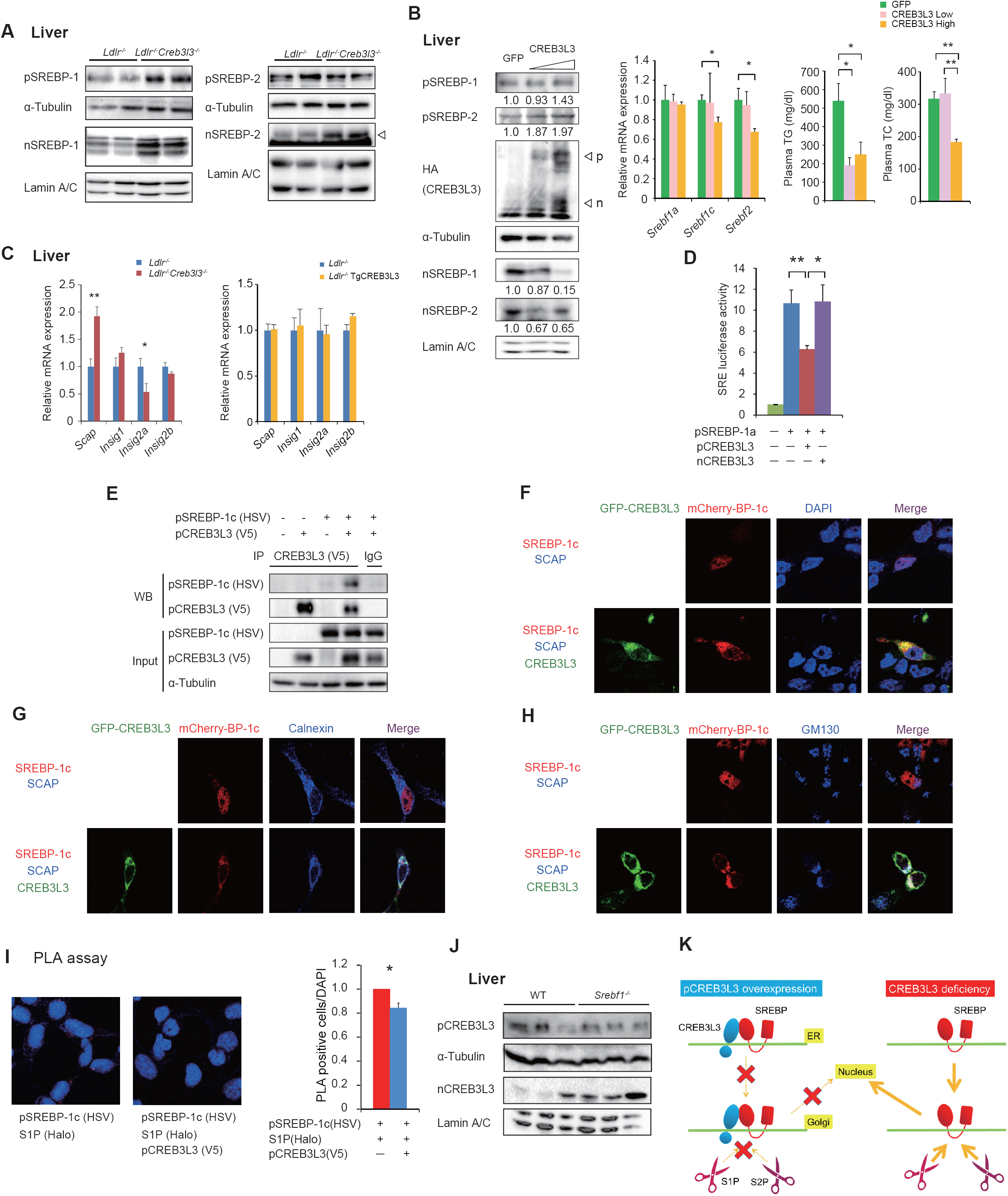
Premature CREB3L3 suppresses the cleavage of SREBPs. (**A**) Immunoblot analysis of SREBPs in hepatic nuclear extracts and total cell lysates from 8-week-old female *Ldlr*^-/-^ and *Ldlr*^-/-^*Creb3l3*^-/-^ mice. Each lane was a pooled sample from three mouse livers. (**B**) Immunoblot analysis of SREBPs in hepatic nuclear extracts and total cell lysates (Each lane was a pooled sample from three mouse livers. The values were indicated as fold changes of band intensity vs GFP infection.), hepatic expression of *Srebfs*, and plasma TG and TC levels in 10-week-old female *Ldlr*^-/-^ and *Ldlr*^-/-^*Creb3l3*^-/-^ mice infected with adenoviruses encoding green fluorescent protein and the low and high dose of full-length CREB3L3 (pCREB3L3) after feeding with normal chow for 6 days. *n* = 4–8; \**p* < 0.05 and \*\**p* < 0.01 vs. *Ldlr*^-/-^ mice. (**C**) Hepatic SREBP cleavage modulator gene expression in 8-week-old female *Ldlr*^-/-^and *Ldlr*^-/-^*Creb3l3*^-/-^, and 8-week-old male *Ldlr*^-/-^ and *Ldlr*^-/-^ TgCREB3L3 mice in a fed state with normal chow. *n* = 4–6; \**p* < 0.05 and \*\**p* < 0.01 vs. *Ldlr*^-/-^ mice. (**D**) pCREB3L3 suppresses SREBP transcriptional activity. pCREB3L3 or active form of CREB3L3 (mCREB3L3), and pSREBP expression vectors were co-transfected with an SRE-luc vector into HEK293 cells. Luciferase activity was determined after 48 h. *n* = 4–8; \**p* < 0.05 and \*\**p* < 0.01 vs. Control. (**E**) Physical association between CREB3L3 and SREBPs. Vectors were co-transfected into HEK293 cells; after 24 h, cell lysates were collected and immunoprecipitated with an anti-V5 antibody. Immunoprecipitants were detected with the indicated antibodies. (**F - H**) The localization of SREBP-1c and CREB3L3 in the cellular component. mCherry-pSREBP-1c (mCherry-BP-1c) and SCAP, with/without GFP-pCREB3L3 (GFP-CREB3L3) were co-transfected into HEK293 cells. DAPI for the nucleus (**F**), Calnexin for ER (**G**), and GM130 for Golgi apparatus (**H**) were immunostained. (**I**) CREB3L3 inhibits SREBPs-S1P interaction. Using DuoLink PLA assay, the red dots showed the pSREBP-1c-S1P association. \**p* < 0.05. (**J**) Immunoblot analysis of CREB3L3 in hepatic nuclear extracts and total cell lysates from 16-week-old male WT and *Srebf1*^-/-^ mice. (**K**) Schematic representation of atherosclerosis development via a competitive transport interaction of SREBPs and CREB3L3.

### Antagonism between CREB3L3 and SREBP occurs at trafficking from ER to Golgi

An immunoprecipitation assay, pCREB3L3 was associated with pSREBP-1c (Figure 6E), indicating a direct association of the two precursor proteins. To determine the effects of pCREB3L3 on the cellular localization of SREBPs, mCherry-tagged pSREBP-1c, SCAP, with or without GFP-tagged pCREB3L3 vectors were co-transfected into HEK293 cells. Immunohistochemistry analysis revealed that SREBP-1c was localized in the nucleus merging with nuclear marker (DAPI) when co-transfected with SCAP indicating SCAP enhanced SREBP cleavage and caused its nuclear transfer (Figure 6F). However, when additionally co-transfected with pCREB3L3, SREBP-1c is then colocalized, merging with CREB3L3 at least not in nucleus, supporting that their direct binding leads to the suppression of SREBP cleavage by pCREB3L3 (Figure 6F). Organelle marker co-transfection indicated that the co-localization of SREBP-1 and CREB3L3 occurs at the ER because both proteins and the ER marker (Calnexin) were all merged (Figure 6G). Without CREB3L3, SREBP-1c and SCAP are dissociated in the nucleus and ER, respectively (Figure 6G). Golgi marker (GM130) in SREBP-1c/SCAP transfection showed a partial signal of SREBP-1c merging at Golgi, presumably a remnant of the uncleaved one and the other partial signal of unmerged one presumably cleaved into the nucleus (Figure 6H). SREBP-1c/SCAP/CREB3L3 co-transfection caused only marginal signaling merge of SREBP-1c-CREB3L3 at Golgi (Figure 6H). The data indicate that CREB3L3 inhibited SCAP escote of SREBP-1c to Golgi by forming the complex, which remained in the ER, thus not allowing the entry of SREBP-1c to the Golgi. To further investigate whether pCREB3L3 disturbs the pSREBPs-S1P complex we used an *in situ* microscopy approach using a proximity ligation assay (PLA). HSV-tagged pSREBP-1c and Halo-tagged S1P were co-transfected into HEK293 cells. Complexes between pSREBP-1c and S1P were observed as red dots around the nucleus (Figure 6I). As CREB3L3 inhibited its complex formation, the red dots were decreased significantly in addition to pCREB3L3 transfection (Figure 6I), which indicates that pCREB3L3 inhibited the formation of the complex between pSREBP-1c and S1P. These findings suggest that the physical association of CREB3L3 with SREBPs inhibits the SCAP-mediated transport of SREBP-1 from the ER to the Golgi, the processing by S1P, and consequently SREBP transcriptional activity. To verify vice versa observation: whether CREB3L3 cleavage is conversely inhibited by SREBP, we evaluated nCREB3L3 protein levels in *Srebf1*^*-/-*^ mice. Hepatic nCREB3L3 protein levels in *Srebf1*^*-/-*^ mice were clearly increased compared with WT mice (Figure 6J). These results support that CREB3L3 and SREBP-1 can antagonize each other by the direct mutual interaction at the precursor protein level.

## Discussion

The current study clearly exhibited that CREB3L3 has impacts on atherosclerotic phenotypes so profoundly that gain- and loss-of-function experiments of a single molecule to date have never shown. *Creb3l3* deletion caused both hypertriglyceridemia and hypercholesterolemia and accelerated aortic atheroma formation in *Ldlr*^-/-^ mice fed either a WD or a normal diet, irrespective of gender, and from both hepatic and intestinal origins. Conversely, hepatic nuclear CREB3L3 overexpression strikingly attenuated WD-induced hyperlipidemia and atherosclerosis progression in *Ldlr*^-/-^ mice. As many pieces of literature including ours and the present work confirmed that the primary role of CREB3L3 is regulation of TG metabolism (*Lee et al., 2011, Nakagawa et al., 2016a, Nakagawa et al., 2016b, Nakagawa et al., 2014, Satoh et al., 2020*), TG per se has been questioned as the major atherosclerosis risk factor, highlighting the cholesterol-related or more comprehensive mechanisms for potential mechanisms of anti-atherogenic effects of CREB3L3.

In the process of clarifying the causative mediators for the anti-atherogenic effect of CREB3L3, FGF21, a major hepatic CREB3L3 target gene which regulates both lipid and glucose metabolism, had been a strong candidate because it was reported to ameliorate atherosclerosis and show hepatic activation of SREBP-2 in *Apoe*^-/-^*Fgf21*^-/-^ mice (*Lin et al., 2015, Wu et al., 2014*). Although the current study also showed that some metabolic phenotypes observed in *Ldlr*^-/-^*Creb3l3*^-/-^ mice were attributed to decreases in plasma FGF21 levels, deficiency of *Fgf21* in *Ldlr*^-/-^TgCREB3L3 mice did not cancel atherosclerosis improvement, distracting the hypothesis that FGF21 is the main contributor of anti-atherosclerosis.

The intestine-specific CREB3L3 KO mice also exhibited atheroma formation, which was comparable to liver-specific KO mice, confirming the role of intestinal CREB3L3 in the anti-atherogenic action. Decreased LXR signaling and increased lipid contents were observed in the intestines of *Ldlr*^-/-^*Creb3l3*^-/-^ mice. Intestinal-specific active form of LXR overexpression in *Ldlr*^-/-^ mice on a WD increased fecal neutral sterol excretion and exhibited protection against atherosclerosis (*Lo Sasso et al., 2010*), supporting the hypothesis that intestinal CREB3L3 contributes to cholesterol metabolism via LXR. L*dlr*^-/-^*Creb3l3*^-/-^ mice had higher plasma plant sterol levels than *Ldlr*^-/-^ mice, explaining that the deficiency of *Creb3l3* in the small intestine increases cholesterol absorption. Collectively, the data suggest that both liver- and intestine-specific CREB3L3 deficiency additively promote atherosclerosis.

We showed that *Creb3l3* deficiency increases levels of hepatic nSREBPs and, consequently, of plasma lipids, which led us to speculate and confirm that CREB3L3 physically competes with SREBPs in respect to ER to Golgi trafficking for cleavage by S1P and S2P. It can be interpreted that CREB3L3 deletion leads to primarily decreased TG catabolism by per se and enhanced lipogenesis as a secondary consequence of SREBP-1c activation, leading to a marked accumulation of TG-rich remnant lipoproteins and severe hypertriglyceridemia. In addition, both increased cholesterol absorption due to absence of intestinal CREB3L3 and cholesterol synthesis mediated by SREBP-2 activation in the liver significantly enriched these lipoproteins with cholesterol and played a major role in the production of more atherogenic lipoproteins.

Functional competition between SREBPs and CREB3L3 implicates profound physiological consequences for lipid and energy regulation. CREB3L3 and SREBPs use the same activation process, regulated intramembrane proteolysis, by which transmembrane proteins are cleaved to release cytosolic domains that translocate into the nucleus and thereby regulate gene transcription. CREB3L3 is cleaved by the processing enzymes, S1P and S2P, in the Golgi apparatus in a manner similar to SREBPs cleavage (*Zhang et al., 2006*). Therefore, we initially hypothesized that pCREB3L3 would competitively inhibit SREBP processing. We showed that CREB3L3 and SREBP-1c are physically interacted (Fig. 6A). It is likely to inhibit trafficking of SREBP-CREB3L3 complex to Golgi, although we could not observe the immunoprecipitation assay of CREB3L3-Insig (data not shown). We have not confirmed complex formation of the three molecules and whether Insig is involved in this potential complex. Precise molecular mechanism needs to be clarified in future. Recently, it was reported that CREB3L3 increases *Insig-2a* expression, which in turn suppresses SREBP activation (*Wang et al., 2016*). Consistently, its expression was decreased in the livers of *Ldlr*^-/-^*Creb3l3*^-/-^ mice, but not changed in *Ldlr*^-/-^TgCREB3L3 mice. Additionally, *Ldlr*^-/-^TgCREB3L3 mice (overexpression of nCREB3L3) did not exhibit apparent altered SREBPs and their target gene activation in the liver. Certainly, adenoviral overexpression of pCREB3L3 in the livers of *Ldlr*^-/-^*Creb3l3*^-/-^ mice reduced nSREBP-1 and nSEEBP-2 protein expression (Figure 6B), indicating that in *Ldlr*^-/-^ mice, SREBP cleavage was regulated by existence of pCREB3L3, rather than Insigs. It was reported that CREB3L3 suppresses LXRα-induced *Srebf-1c* expression by inhibiting LXRα binding to *Srebf-1c* promoter (*Min et al., 2013*); however, in our data settings, *Ldlr*^-/-^TgCREB3L3 mice exhibited no changes in *Srebf1c* and its target genes.

We propose the new concept that pCREB3L3 and pSREBPs physically interact at the ER and inhibit transportation of SREBPs to the Golgi apparatus, and even if transferred, competes with SREBPs in access to S1P in the Golgi apparatus. Loss of this interaction due to CREB3L3 deficiency induces SREBP-1 and −2 cleavage and promotes TG and cholesterol synthesis induction (Figure 6K). This mechanism also explains why the overexpression of nuclear CREB3L3 did not suppress hepatic SREBP target genes (Figure 5B), because nuclear CREB3L3 does not compete with pSREBPs. In the normal liver, *Creb3l3* is upregulated during fasting and conversely, downregulated under fed conditions, while *Srebf1c* is regulated in the reciprocal way. Thus, the encounter of the two factors does not actively occur on the ER under normal nutritional states. However, in metabolic disturbances with high atherogenic risks, such as *db/db* or *ob/ob* mice, two factors could be expressed collaterally (*Nakagawa et al., 2014*) and interact with and inhibit each other. Finally, enhancement of nCREB3L3 with decreased pCREB3L3 in *Srebf1*^-/-^ liver (Figure 6J) supports this hypothesis. Functional antagonism between CREB3L3 and SREBPs is consistent in atherosclerosis based upon the anti-atherogenic action of CREB3L3 from the current data and pro-atherogenic action of SREBP-1 from our previous work (*Karasawa et al., 2011*). CREB3L3 and SREBPs are regulators of catabolism and anabolism of lipids, respectively, it is conceivable to configure a mechanism by which mutual interaction and balance of the counterpartners maintain the whole-body energy balance and atherosclerosis risks.

Collectively, our study is the first to identify the crucial role of CREB3L3 enterohepatic interplay in lipid metabolism and atherosclerosis prevention. CREB3L3 could be a new atherosclerosis target. Protection from atherosclerosis by overexpressed nuclear CREB3L3 mice was more than expected from the amelioration of hyperlipidemia in *Ldlr*^-/-^TgCREB3L3 mice. CREB3L3 is deeply involved in cellular stress and inflammation, which we have not investigated in the current study. Therefore, further study is warranted in these aspects.

## Materials and Methods

### Mice

This project was approved and conducted under the guidelines of the Animal Care Committee of the University of Tsukuba. *Creb3l3*^*tm1*.*1Sad/J*^ *(Creb3l3*^-/-^) mice (*Luebke-Wheeler et al., 2008*) and *Ldlr*^-/-^ mice (*Ishibashi et al., 1993*) were purchased from Jackson Laboratory (Bar Harbor, ME, USA). *Creb3l3*^-/-^ mice were crossed onto an *Ldlr*^-/-^ background to generate *Ldlr*^-/-^*Creb3l3*^-/-^ mice. Transgenic mice overexpressing amino acids 1–320 of human CREB3L3 under control of the phosphoenolpyruvate carboxykinase promoter on the C57BL/6J background (hereafter referred to as TgCREB3L3) were generated as previously described (*Nakagawa et al., 2014*). *Fgf21*^-/-^ mice were kindly provided by Prof. Morichika Konishi and Nobuyuki Ito (*Hotta et al., 2009*). TgCREB3L3 mice were crossed with *Ldlr*^-/-^ mice to produce *Ldlr*^-/-^TgCREB3L3 mice and then crossed with *Fgf21*^-/-^ mice to produce *Ldlr*^-/-^*Fgf21*^-/-^TgCREB3L3 mice. *Creb3l3*^*flox/flox*^ (flox) mice were generated using the CRISPR/Cas 9 system as previously described (*Nakagawa et al., 2016a*). Flox mice were crossed with B6.Cg-Tg(Alb-Cre)21Mgn/J (albumin Cre Tg, from Jackson Lab) (*Yakar et al., 1999*) and/or villin Cre Tg mice (from Jackson Lab) (*Madison et al., 2002*) to produce liver-specific CREB3L3 knockout (LKO), small intestine-specific CREB3L3 knockout (IKO), and both liver- and small intestine-specific CREB3L3 knockout (DKO) mice. These mice were crossed with *Ldlr*^-/-^ mice generating *Ldlr*^-/-^flox, *Ldlr*^-/-^LKO, *Ldlr*^-/-^IKO, and *Ldlr*^-/-^DKO mice, respectively. *Srebf1*^-/-^ mice were generated as previously described (*Shimano et al., 1997*). Sixteen-week-old male WT and *Srebf1*^-/-^ mice were fasted for 24 hrs and then fed with high-sucrose diet for 12 hrs (*Nakagawa et al., 2006*). All mice were maintained on normal chow (Oriental Yeast Company, Tokyo, Japan) and a 14-h light/10-h dark cycle. For atherosclerosis analyses, mice were fed WD (D12079B [34% sucrose, 21% fat, 0.15% cholesterol], Research Diet) under the indicated terms (*Karasawa et al., 2011*). For adenoviral infection, 8- to 10-week-old female *Ldlr*^-/-^*Creb3l3*^-/-^ mice were infected with the indicated adenovirus at 1.0 (low) and 5.0 (high dose) × 10^8^ pfu/g body weight (BW); samples were collected 6 days later while in a fed state. All animal husbandry procedures and animal experiments were consistent with the University of Tsukuba Regulations of Animal Experiment and were approved by the Animal Experiment Committee of the University of Tsukuba.

### Determination of metabolic parameters

Plasma levels of glucose, triglycerides (TG), TC, FFA, alanine aminotransferase (ALT), and aspartate aminotransferase (AST) were measured using Wako enzymatic kits (Wako Pure Chemical Industries). Plasma insulin was measured with a mouse insulin enzyme-linked immunosorbent assay (ELISA) kit (Sibayagi). Plasma FGF21 was measured with a mouse/rat FGF21 Quantikine ELISA kit (R&D Systems). Hepatic TG and TC contents were measured as previously described(*Nakagawa et al., 2006*). Intestinal TG and TC contents were measured using the same protocol. Plasma ApoA4 was detected by western blotting with an anti-ApoA4 antibody (sc-19036; Santa Cruz Biotechnology).

### HPLC analysis

For lipoprotein distribution analysis, pooled plasma samples from four to five mice per group were analyzed via upgraded HPLC analysis as previously described (Skylight Biotech Inc.) (*Okazaki et al., 2005*).

### Isolation of VLDL fraction

VLDL (*d* < 1.006 g/ml) was isolated via ultracentrifugation with a TLA120.2 rotor (Beckman Coulter). VLDL fractions were separated by sodium dodecyl sulfate (SDS)-polyacrylamide gel electrophoresis (PAGE) and subjected to Coomassie brilliant blue staining.

### Determination of plasma sterol levels

For sterol distribution analysis, pooled plasma samples were quantified using a gas chromatography method (Skylight Biotech).

### Immunoblotting

Total cell and nuclear fraction lysates were prepared as previously described(*Ide et al., 2004*), separated by SDS-PAGE, and subjected to western blot analysis using antibodies against SREBP-1 (sc-12332; Santa Cruz), SREBP-2 (10007663; Cayman Chemical), α-Tubulin (05-829; Millipore), Lamin A/C (#2032, Cell Signaling Technologies), V5 (R960; Life Technologies), HSV (69171-3; Novagen), and HA (3F10) (Roche) antibodies.

### Immunoprecipitation

HEK293 cells were maintained in Dulbecco’s modified Eagle’s medium supplemented with 10% fetal bovine serum and penicillin/streptomycin. Indicated plasmids were transfected with X-tremeGENE 9 (Roche) according to the manufacturer’s instructions. V5-tagged full-length mouse *Creb3l3* cDNA was inserted into pcDNA3.1 (Invitrogen); GFP-tagged full-length mouse *Creb3l3* cDNA was inserted into pEGFP (GFP-CREB3L3) (Clontech), mCherry-tagged human SREBP-1c was inserted into pmCherry (mCherry-SREBP-1c) (Clontech), and HA-tagged hamster SCAP, HSV-tagged human SREBP-1c and HSV-tagged human SREBP-2 were inserted into pCMV. Cell lysates were immunoprecipitated with antibodies against V5 and immunoprecipitants were subjected to immunoblotting with the indicated antibodies as previously described (*Nakagawa et al., 2014*).

### Duolink proximity ligation (PLA) assay

HEK293 cells were co-transfected with HSV-tagged pSREBP-1c and Halo-tagged S1P with/without pCREB3L3. Cells were fixed with 3.7% formalin in PBS for 30 min before being permeabilized with 0.2% Triton X-100 in PBS for 10 min. Cells were then subjected to the PLA assay using the Duolink red kit (Sigma-Aldrich) according to the manufacturer’s instructions.

### Immunocytochemistry

HEK293 cells were transfected with mCherry-tagged pSREBP-1c, SCAP, and GFP-tagged pCREB3L3 using X-tremeGENE 9 (Roche). Cells were grown on coverslips, fixed with 4 % paraformaldehyde for 15min, and permeabilized with 0.1 % Triton X-100 for 5min. After blocking in 1% BSA for 30min, the cells were incubated with primary and secondary antibodies for 1h each. The ER and Golgi apparatus were stained using anti-Calnexin (610523, BD Biosciences) and anti-GM130 antibodies (610822, BD Biosciences), respectively. Immunoreactive complexes were visualized with Alexa Fluor 405-conjugated secondary antibody (ab175660, Abcam) and nuclei were visualized by staining with 4’,6-diamidino-2-phenylindole.

### Promoter analysis

HEK293 cells were transfected with the indicated luciferase reporter, expression plasmids, and a reference pRL-SV40 plasmid (Promega) using X-tremeGENE 9 (Roche). SRE-luc vector (*Ide et al., 2004, Ide et al., 2003, Yoshikawa et al., 2003*) and human SREBP-1a^41,42^ were described previously. After a 48-h incubation, firefly luciferase activity in cells was measured and normalized to *Renilla* luciferase activity.

### Atherosclerotic lesion analysis

Ten ∼ eleven-week-old male and female *Ldlr*^-/-^ and *Ldlr*^-/-^*Creb3l3*^-/-^ mice were fed a WD containing 34% sucrose, 21% fat, and 0.15% cholesterol (D12079B; Research Diet) for 5 weeks and 3 months. Ten–eleven-week-old male *Ldlr*^-/-^ and *Ldlr*^-/-^TgCREB3L3 mice were fed a WD for 3 months. Mice were subsequently euthanized to extract their hearts and aortas. Hearts were fixed in 4% formalin for >48 h. The basal half of each heart was embedded in Tissue-Tek OCT compound (Sakura Finetek). Cross-sections were stained with Oil Red O and hematoxylin. Aortas were cut along the midline from the iliac arteries to the aortic root, pinned flat, and treated with Sudan IV for 15 min to stain lesions, followed by 70% ethanol destaining and fixation in 4% phosphate-buffered formalin(*Karasawa et al., 2011*). Atherosclerotic lesions were quantified using Photoshop CS software (Adobe Systems Inc.).

### TG production

Mice were deprived of food for 24 h and subsequently injected with Triton WR-1339 (0.5 mg/g BW; Sigma-Aldrich) via the tail veins to block the clearance of nascent ApoB-containing lipoproteins. Blood samples were collected at 0, 30, 60, and 120 min post-injection (*Karasawa et al., 2011*).

### Postprandial TG response

Mice were deprived of food for 16 h, followed by oral administration of 200 μl of olive oil(*Lee et al., 2011*). Blood samples were collected at 0, 3, 6, and 9 h post-administration.

### Intestinal TG absorption

Mice were fasted for 3 h and injected with Triton WR-1339 (1 mg/g BW, Sigma-Aldrich) via the tail veins. After 2 h injection, mice were administrated with 100 μl of Olive Oil orally (*Uchida et al., 2012*). Blood samples were collected for up to 3 h after the injection.

### Determination of plasma LPL activity

Mice were injected with 100 U/kg BW of heparin (Novo Heparin, Mochida Pharmaceutical Co., Ltd) via the tail veins. Blood samples were collected at 20 min post-administration. Plasma LPL activity was determined using an LPL activity assay kit (Roar Biochemical, Inc) according to the manufacturer’s instructions.

### Preparation of recombinant adenovirus

cDNAs encoding human full-length of CREB3L3 (NM_032607.2) and green fluorescent protein were cloned into pENTR4 vectors (Life Technologies). In addition, adenovirus vectors were recombined with pAd/CMV/V5-DEST vectors (Life Technologies). Recombinant adenoviruses were produced in 293A cells (Invitrogen) and purified via CsCl gradient centrifugation, as previously described (*Nakagawa et al., 2006*).

### Analysis of gene expression

Total RNA was isolated from cells and tissues using Trizol reagent (Invitrogen) and Sepasol (Nacalai). Real-time PCR analysis templates were prepared via cDNA synthesis (Invitrogen) from total RNA. Real-time PCR was performed using the ABI Prism 7300 System (Applied Biosystems, Inc) with SYBR Green Master Mix (Roche) and TB Green Premix EX Taq II (TAKARA Bio) (*Fujimoto et al., 2013*). Primer sequences are described in Supplementary Table 1.

### Statistical analyses

Statistical significance was determined using unpaired Student’s *t*-tests and one-way ANOVA with Tukey’s post hoc using the GraphPad Prism software. Differences with *p* values <0.05 were considered significant. Data are expressed as means ± standard errors of the mean

## Acknowledgements

The authors would like to thank Enago (www.enago.jp) for the English language review. This work was supported by Grants-in-Aid from the Ministry of Science, Education, Culture and Technology of Japan (25282214 and 16H03253, to Y.N., and 17H06395 and 18H04051 to H.Shimano), AMED-CREST (to Y.N. and H.Shimano), Japan Heart Foundation/Novartis Grant for Research Award on Molecular and Cellular Cardiology (to Y.N.), Uehara Memorial Foundation (to Y.N.), Ono Medical Research Foundation (to Y.N.), Mochida Memorial Foundation for Medical and Pharmaceutical Research (to Y.N.), Suzuken Memorial Foundation (to Y.N.), Senshin Medical Research Foundation (to Y.N.), Takeda Science Foundation (to Y.N.), Japan Foundation for Applied Enzymology (to Y.N.), Banyu Life Science Foundation International (to Y.N.), and Yamaguchi Endocrine Research Foundation (to Y.N.).

## Author contributions

Y.N. and H.Shimano designed the experiments and wrote the manuscript. Y.N., Y.W., S-I.H., K.O., A.O., Y.Y., K.K., H.O., Y.Mizunoe, and M.A. performed the experiments. M.K. and N.I. generated *Fgf21*^*-/-*^ mice. Y.O., Y.Murayama, H.I., T.M., and H.Sone were involved in project planning. N.Yamada supervised this study and contributed crucial ideas to the project.

## Supplement figure legend

**Figure 1—figure supplement 1.**
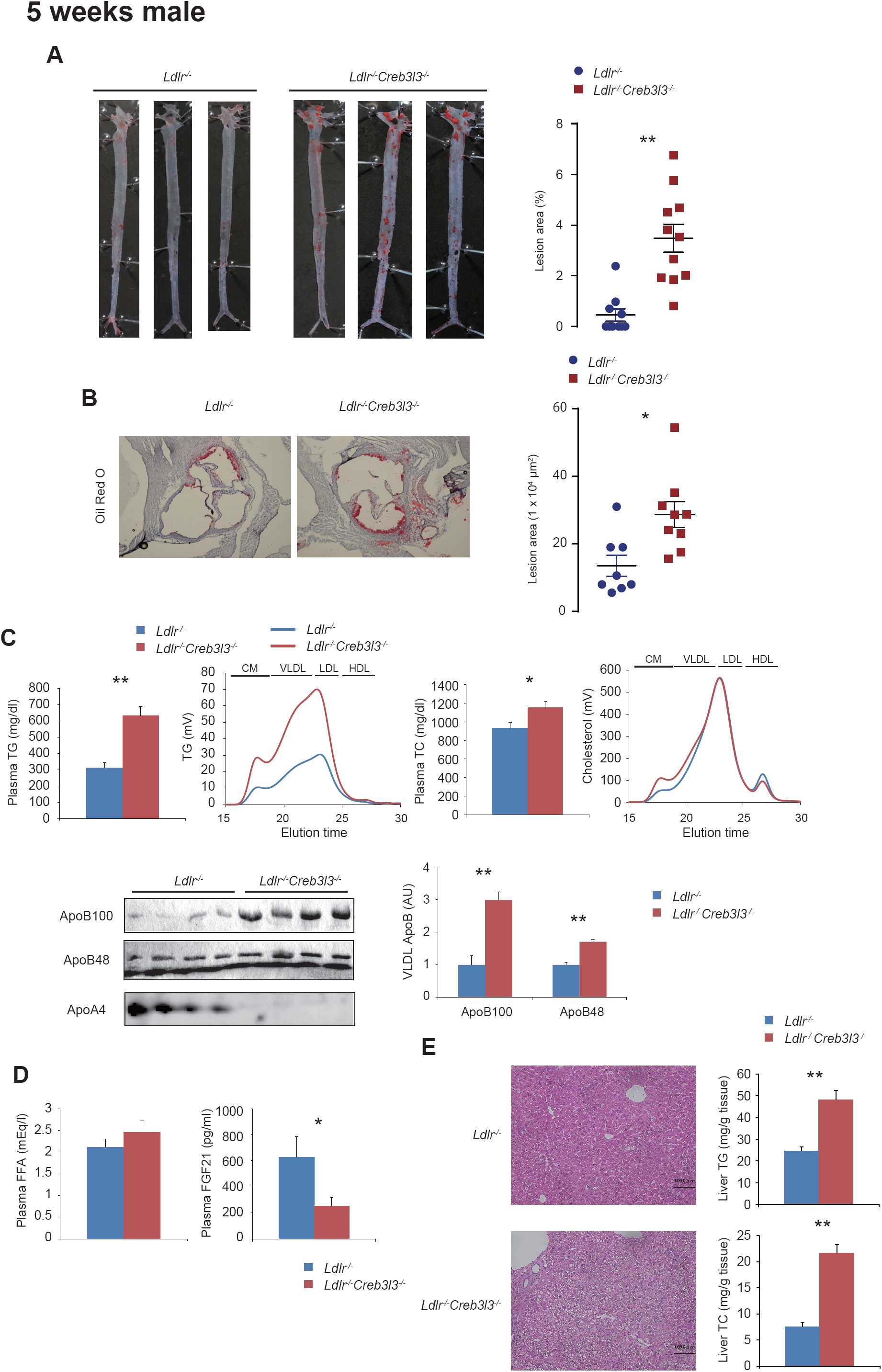
Male *Ldlr*^-/-^ *Creb3l3*^-/-^ mice promote atherosclerosis feeding a WD for 5 weeks. Ten ∼ eleven-week-old male *Ldlr*^-/-^ and *Ldlr*^-/-^*Creb3l3*^-/-^ mice were fed a WD for 5 weeks; samples were collected in a fed state. (**A**) Representative images of entire Sudan IV-stained aortas from *Ldlr*^-/-^ (*n* = 11) and *Ldlr*^-/-^*Creb3l3*^-/-^ (*n* = 11) mice. The surface area occupied by lesions was quantified. \*\**p* < 0.01 vs. *Ldlr*^-/-^ mice. (**B**) Representative aortic root sections from *Ldlr*^-/-^ (*n* = 11) and *Ldlr*^-/-^*Creb3l3*^-/-^ (*n* = 11) mice. Cross-sections were stained with Oil Red O and hematoxylin. Aortic root lesion areas were quantified. \**p* < 0.05 vs. *Ldlr*^-/-^ mice. (**C**) Plasma TG and TC levels, *n* = 11 each; HPLC analysis of plasma lipoprotein profiles specific for plasma TG and cholesterol. ApoB100 and ApoB48 were isolated from VLDL fractions via ultracentrifugation, subjected to SDS-PAGE, stained with CBB, and quantified. *n* = 6–7 per group; \**p* < 0.05 and \*\**p* < 0.01 vs. *Ldlr*^-/-^ mice. Plasma ApoA4 levels were determined by WB. (**D**) Plasma FFA and FGF21 levels. *n* = 11 each; \**p* < 0.05 vs. *Ldlr*^-/-^ mice. (**E**) Histology of liver sections, and liver TG and TC levels, *n* = 12–15 per group; \**p* < 0.05 vs. *Ldlr*^-/-^ mice.

**Figure 1—figure supplement 2.**
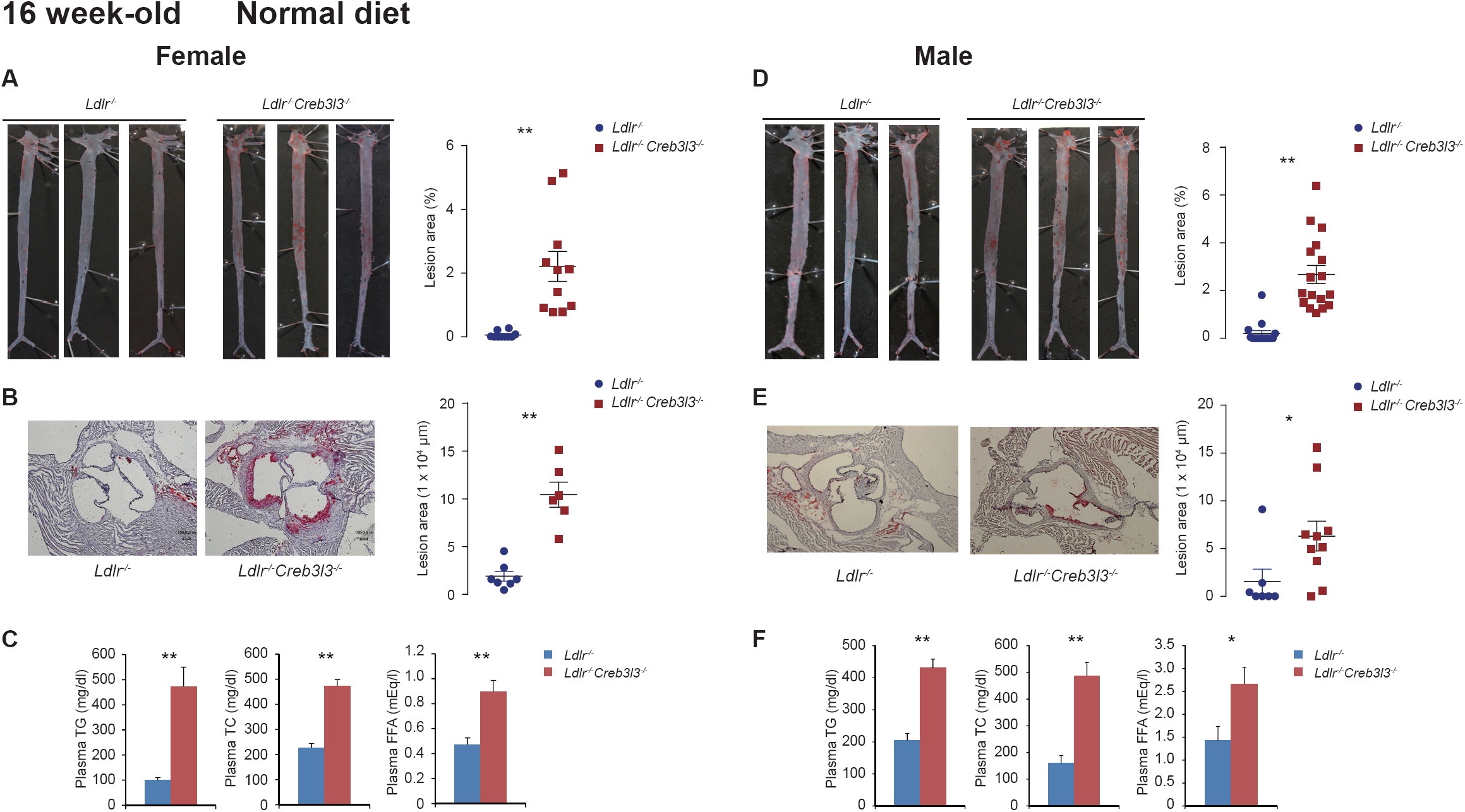
*Ldlr*^-/-^*Creb3l3*^-/-^ mice promote atherosclerosis even feeding with normal chow. Sixteen-week-old female (**A**–**C**) and male (**D–F**) *Ldlr*^-/-^ and *Ldlr*^-/-^*Creb3l3*^-/-^ mice were fed a normal diet. Samples were collected in a fed state. (**A**) Representative aortic root sections from female *Ldlr*^-/-^ (*n* = 10) and *Ldlr*^-/-^*Creb3l3*^-/-^ (*n* = 11) mice. Cross-sections were stained with Oil Red O and hematoxylin. Aortic lesion areas were quantified. \*\**p* < 0.01 vs. *Ldlr*^-/-^ mice. (**B**) Representative images of entire Sudan IV-stained aortas from *Ldlr*^-/-^ (*n* = 7) and *Ldlr*^-/-^*Creb3l3*^-/-^ (*n* = 6) mice. The surface area occupied by lesions was quantified. \*\**p* < 0.01 vs. *Ldlr*^-/-^ mice. (**C**) Plasma TG, TC, and FFA levels of female *Ldlr*^-/-^ (*n* = 6) and *Ldlr*^-/-^*Creb3l3*^-/-^ (*n* = 7) mice. \*\**p* < 0.01 vs. *Ldlr*^-/-^ mice. (**D**) Representative aortic root sections from male *Ldlr*^-/-^ (*n* = 15) and *Ldlr*^-/-^*Creb3l3*^-/-^ (*n* = 17) mice. Cross-sections were stained with Oil Red O and hematoxylin. Aortic lesion areas were quantified. \*\**p* < 0.01 vs. *Ldlr*^-/-^ mice. (**E**) Representative images of entire Sudan IV-stained aortas from male *Ldlr*^-/-^ (*n* = 7) and *Ldlr*^-/-^*Creb3l3*^-/-^ (*n* = 10) mice. The surface area occupied by lesions was quantified. \**p* < 0.05 vs. *Ldlr*^-/-^ mice. (**F**) Plasma TG, TC, and FFA levels of male *Ldlr*^-/-^ (*n* = 10) and *Ldlr*^-/-^*Creb3l3*^-/-^ (*n* = 14) mice. \**p* < 0.05 and \*\**p* < 0.01 vs. *Ldlr*^-/-^ mice.

**Figure 1—figure supplement 3.**
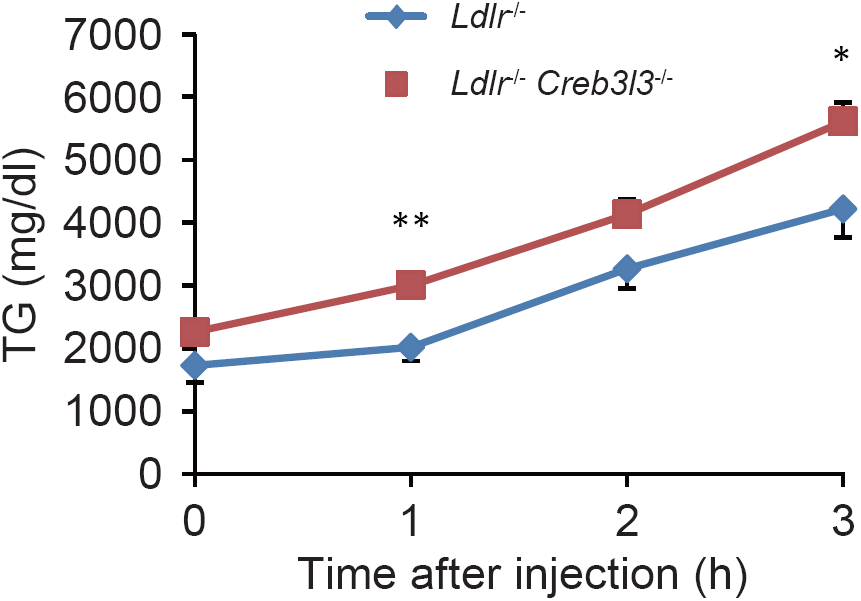
*Ldlr*^-/-^ *Creb3l3*^-/-^ mice increase intestinal TG absorption. Eight-week-old female *Ldlr*^-/-^ and *Ldlr*^-/-^*Creb3l3*^-/-^ mice were fasted for 3 h and intravenously injected with Triton WR-1339. After 2 h of injection, mice were orally administrated 100 μl of olive oil. Plasma was collected at 0, 1, 2, and 3 h after administration. n = 7 each, * p < 0.05, ** p < 0.01, vs *Ldlr*^-/-^ mice.

**Figure 3—figure supplement 1.**
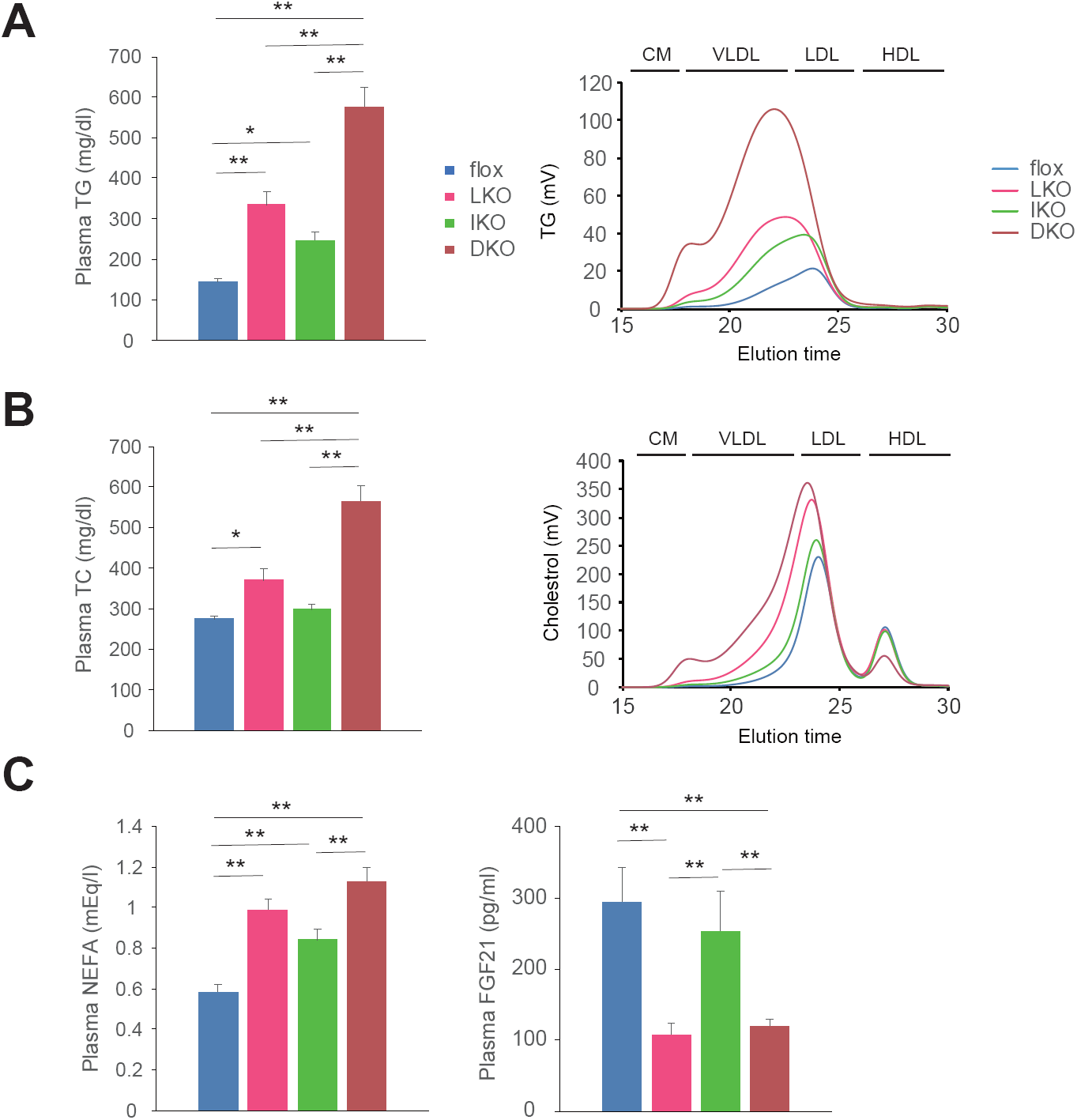
Plasma lipoprotein profiles in female tissue-specific *Creb3l3* deletion in *Ldlr*^-/-^ mice. Samples were collected from 8-week-old female *Ldlr*^-/-^ flox, *Ldlr*^-/-^ LKO, *Ldlr*^-/-^ IKO, and *Ldlr*^-/-^ DKO mice in a fed state. (**A, B**) Plasma TG (**A**), TC (**B**) levels. *n* = 17–29 per group; \**p* < 0.05 and \*\**p* < 0.01 among genotypes. HPLC analysis of plasma lipoprotein profiles of TG and cholesterol. (**C**) Plasma NEFA (*n* = 18–25 per group) and FGF21 (*n* = 16–17 per group) levels. \*\**p* < 0.01 among genotypes.

**Figure 4—figure supplement 1.**
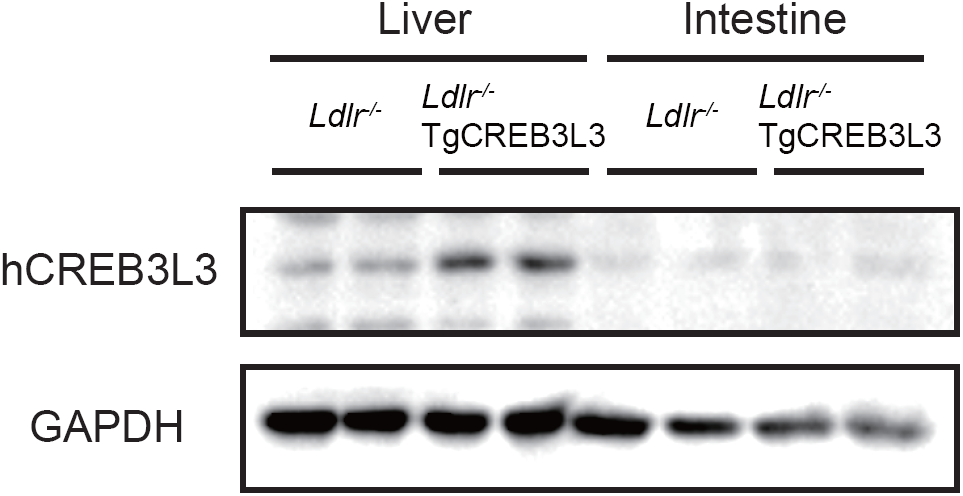
Ectopic active form of CREB3L3 protein in liver and intestine of *Ldlr*^-/-^TgCREB3L3 mice. The ectopic active form of CREB3L3 protein levels in the liver and small intestine of 8-week-old male *Ldlr*^-/-^ and *Ldlr*^-/-^TgCREB3L3 mice were determined by western blotting.

**Figure 5—figure supplement 1.**
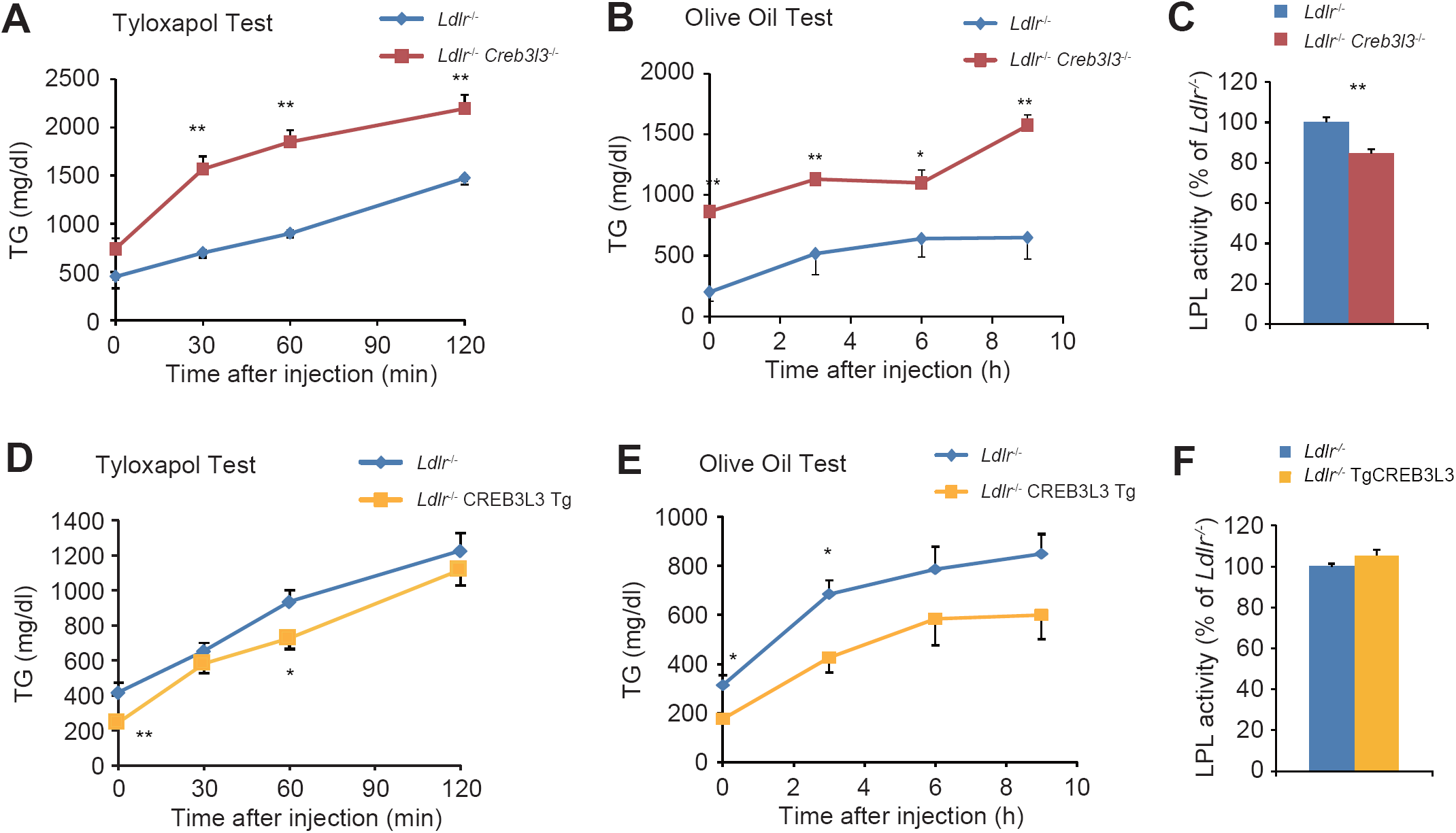
Very low-density lipoprotein secretion and TG clearance in *Ldlr*^-/-^*Creb3l3*^-/-^ and *Ldlr*^-/-^TgCREB3L3 mice. (**A, D**): TG production rates (Tyloxapol test) in 8-week-old female *Ldlr*^-/-^ (*n* = 6) and *Ldlr*^-/-^*Creb3l3*^-/-^ (*n* = 11) mice (**A**) and male *Ldlr*^-/-^ (*n* = 6) and *Ldlr*^-/-^TgCREB3L3 (*n* = 6) mice (**D**). Mice were starved for 24 h and intravenously injected with Triton WR-1339. Plasma was collected at 0, 30, 60, and 120 min post-injection. \*\**p* < 0.01 vs. *Ldlr*^-/-^ mice. (**B, E**) Postprandial TG responses (Olive oil test) in 9-week-old female *Ldlr*^-/-^ (*n* = 7–8) and *Ldlr*^-/-^*Creb3l3*^-/-^ (*n* = 8) mice (**B**) and male *Ldlr*^-/-^ (*n* = 7) and *Ldlr*^-/-^TgCREB3L3 (*n* = 5) mice (**E**). Mice were starved for 16 h, followed by oral administration of 200 μl of olive oil. Plasma was collected at 0, 3, 6, and 9 h post-administration. \**p* < 0.05 and \*\**p* < 0.01 vs. *Ldlr*^-/-^ mice. (**C, F**) Plasma lipoprotein lipase activity in 8-week-old female *Ldlr*^-/-^ (*n* = 5) and *Ldlr*^-/-^*Creb3l3*^-/-^ mice (*n* = 6) (**C**) and male *Ldlr*^-/-^ (*n* = 5) and *Ldlr*^-/-^TgCREB3L3 (*n* = 5) mice (**F**). \**p* < 0.05 vs. *Ldlr*^-/-^ mice.

**Supplementary Table 1.**
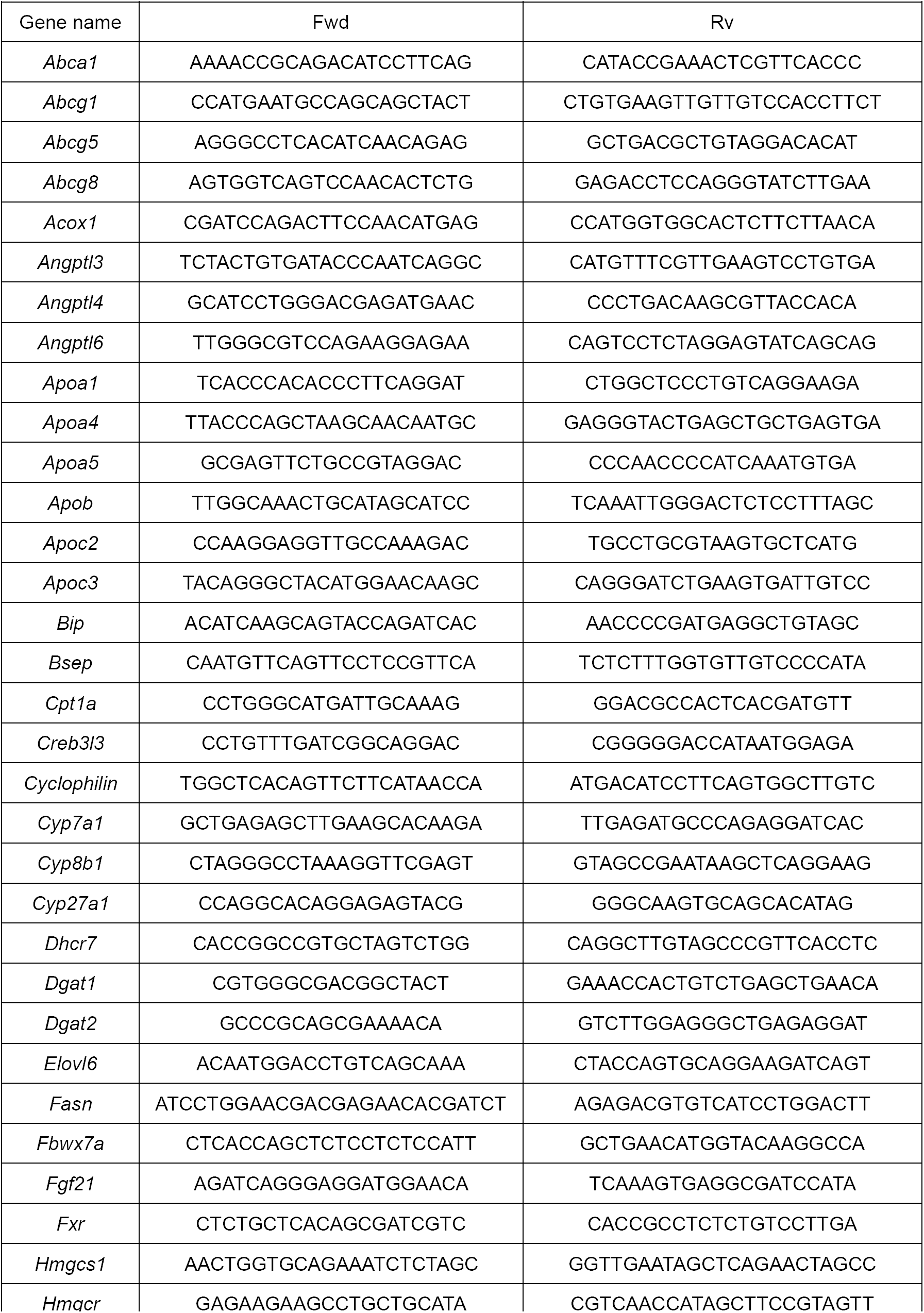
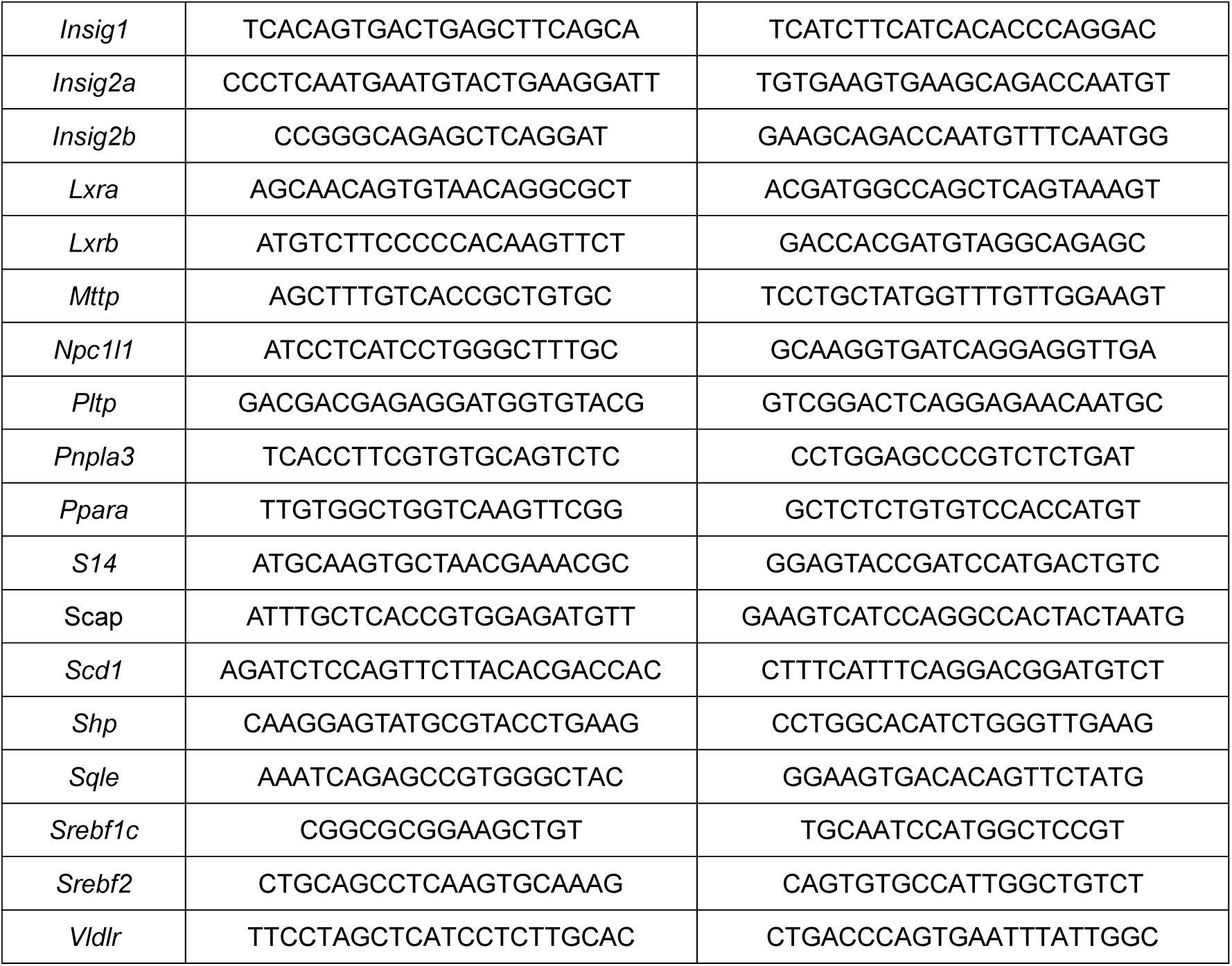
Primers used for real-time PCR analysis.

